# Plexin-A2 Promotes The Proliferation And The Development Of Tumors From Glioblastoma Derived Cells

**DOI:** 10.1101/2021.07.12.452132

**Authors:** Shira Toledano, Adi D. Sabag, Tanya liburkin-Dan, Ofra Kessler, Gera Neufeld

**Affiliations:** Cancer research center, The Bruce Rappaport Faculty of Medicine, Technion, Israel Institute of Technology, Haifa, Israel; Division of Allergy & Clinical Immunology, Bnai-Zion medical Center, Haifa, Israel

**Author notes:** To whom correspondence should be addressed: Dr. Gera Neufeld.

## Abstract

The semaphorin guidance factors receptor Plexin-A2 transduces sema6A and sema6B signals and when associated with neuropilins can also transduce sema3C signals. Inhibition of plexin-A2 expression in U87MG glioblastoma cells resulted in strong inhibition of cell proliferation and tumor forming ability. Knock-out of the plexin-A2 gene using CRISPR/Cas9 also inhibited cell proliferation which was rescued following re-expression of the plexin-A2 cDNA or expression of a truncated plexin-A2 lacking the extracellular domain. Inhibition of plexin-A2 expression resulted in cell cycle arrest at the G2/M stage, and was accompanied by changes in cytoskeletal organization, cell flattening, and by the expression of senescence associated β-galactosidase. It was also associated with reduced AKT phosphorylation and enhanced phosphorylation of p38MAPK. We find that the pro-proliferative effects of plexin-A2 are mediated by FARP2 and FYN since mutations in the FARP2 binding domain of plexin-A2 or in the FYN phosphorylation sites of plexin-A2 compromised the rescue of the proliferative activity by the plexin-A2 intracellular domain. Our results suggest that plexin-A2 may represent a novel target for the development of anti-tumorigenic therapeutics.

## INTRODUCTION

The semaphorins were initially characterized as axon guidance factors, but have subsequently been found to regulate angiogenesis, lymphangiogenesis and immune responses and to modulate tumor progression (Gu and Giraudo, 2013;Eissler and Rolny, 2013;Neufeld *et al*., 2016). The nine receptors of the plexin family function as semaphorin receptors (Tamagnone *et al*., 1999;Takahashi *et al*., 1999) and are subdivided into four type-A, three type-B and single type-C and type-D receptors. They are single pass membrane receptors that are characterized by the presence of a GTPase activating (GAP) domain in their intracellular domain (Hota and Buck, 2012;Kong *et al*., 2016). Upon activation by semaphorins they induce the local collapse of the cytoskeleton in target cells and thus function mostly as repulsive guidance factors. Most semaphorins bind directly to plexins except for six of the seven class-3 semaphorins which bind to receptors of the neuropilin family. Some semaphorins such as sema4D induce tumor progression and tumor angiogenesis following their binding to class-B plexin receptors (Basile *et al*., 2004;Conrotto *et al*., 2005) Neuropilins do not transduce semaphorin signals on their own due to their short intracellular domains. Following the binding of the class-3 semaphorins, they associate with type-A plexins or with plexin-D1, thereby activating plexin mediated signal transduction (Janssen *et al*., 2012;Neufeld *et al*., 2016). Several class-3 semaphorins such as sema3A, sema3B and sema3F have been found to function as potent inhibitors of tumor angiogenesis and consequently as inhibitors of tumor progression (Kessler *et al*., 2004;Bielenberg *et al*., 2004;Varshavsky *et al*., 2008;Casazza *et al*., 2010;Casazza *et al*., 2011;Casazza *et al*., 2012;Mumblat *et al*., 2015;Yang *et al*., 2015). Sema3C and sema3F also inhibit tumor lymphangiogenesis and inhibit tumor metastasis mediated by lymph vessels (Bielenberg *et al*., 2004;Mumblat *et al*., 2015;Doci *et al*., 2015). Notably, sema3C can also function as a promoter of tumor progression as it also binds to the plexin-B1 receptor (Peacock *et al*., 2018).

Sema3C binds to the neuropilin-1 and to the neuropilin-2 receptors. Upon stimulation with sema3C these receptors associate with the plexin-D1 receptor which functions as the primary transducer of sema3C signals (Gitler *et al*., 2004;Smolkin *et al*., 2018). In addition, neuropilin-1 and neuropilin-2 can also associate with the plexin-A2 receptor to transduce sema3B and sema3C signals (Toyofuku *et al*., 2008;Sabag *et al*., 2014;Smolkin *et al*., 2018;Yin *et al*., 2021). Plexin-A2 is one of the four type-A plexins. It functions as a direct binding and signal transducing receptor for sema6A and sema6B (Tawarayama *et al*., 2010). When expressed at high non-physiological concentrations plexin-A2 in complex with neuropilin-1 was also found to convey signals induced by sema3A and possibly also by additional class-3 semaphorins (Janssen *et al*., 2012;Sabag *et al*., 2014). There is some evidence suggesting that high plexin-A2 expression levels are linked to worse prognosis in glioblastoma multiforme (Oncomine, Murat brain dataset, (Fig. S1A)) (Murat *et al*., 2008;Man *et al*., 2014), and in prostate cancer (Tian *et al*., 2013). In glioma stem cells sema3C was found activate Rac1/nuclear factor (NF)-kappaB signaling via plexin-A2/plexin-D1 to promote the survival of these cancer stem cells (Man *et al*., 2014). However, in a recent report these same authors found that plexin-A2 does not convey sema3C signals (Christie *et al*., 2021). They also found that plexin-A2 associates stably with the plexin-A4 receptor which also functions, like plexin-A2, as a receptor for sema6A and sema6B (Suto *et al*., 2005;Haklai-Topper *et al*., 2010;Christie *et al*., 2021). Interestingly, silencing sema6B or plexin-A4 expression in human umbilical vein derived endothelial cells strongly inhibits their proliferation suggesting that plexin-A4 mediated autocrine sema6B signaling promotes their survival/proliferation and inhibits tumor development from glioblastoma derived cells (Kigel *et al*., 2011). Lastly, plexin-A2 also functions as a receptor for KIAA1199, an oncogenic protein that transmits pro-survival and invasiveness promoting signals by stabilization of the epidermal growth factor receptor thus protecting against sema3A induced apoptosis in keratinocytes (Shostak *et al*., 2014), suggesting that KIAA1199 may also be involved in the transduction of plexin-A2 mediated pro-proliferative signals in glioblastomas.

To better understand the involvement of plexin-A2 in glioblastomas we have inhibited the expression of plexin-A2 in U87MG glioblastoma multiforme derived cells using several methods including gene knock-out with CRISPR/Cas9. This resulted in an almost complete inhibition of their proliferation and their tumor forming ability and was also accompanied by profound changes in their cytoskeletal organization that resembled changes associated with cellular senescence. Similar changes were observed following the silencing of plexin-A2 expression in A172 glioblastoma multiforme derived cells. In contrast, silencing plexin-A2 expression had no effect in glioblastoma derived cells containing mutations in the p53 gene. The inhibition of cell proliferation as well as the cytoskeletal changes could be rescued by the expression of cDNAs encoding either full length plexin-A2 or a truncated plexin-A2 lacking the entire extracellular domain. We present evidence suggesting that the pro-proliferative activity of plexin-A2 is not dependent on autocrine activation by sema3C and further show evidence suggesting that the pro proliferative activities of plexin-A2 also require the plexin-A4 receptor and are mediated, at least in part, by the secondary messengers FARP2, FYN, AKT and p38MAPK. Our results suggest that targeting plexin-A2, primarily in glioblastomas that retain wild type p53 expression, may turn out to have therapeutic value.

## RESULTS

### Silencing plexin-A2 expression in U87MG cells inhibits their proliferation and their tumor forming ability

U87MG glioblastoma cells express the four class-A plexins as well as both neuropilin receptors but not the plexin-D1 receptor (Kigel *et al*., 2011;Sabag *et al*., 2014;Smolkin *et al*., 2018). We have silenced the expression of plexin-A2 in U87MG and A172 glioblastoma cells as well as in human umbilical vein derived endothelial cells (HUVEC) using several specific shRNA species. The silencing resulted in strong inhibition of cell proliferation (Figs. 1A-C & S1B). When U87MG cells in which the expression of plexin-A2 was silenced were implemented subcutaneously in immune deficient mice, the development of tumors from the silenced cells was also very strongly inhibited (Figs. 1D & 1E). Both the U87MG and A172 cells express wild type p53. Interestingly, Silencing the expression of plexin-A2 in glioblastoma multiforme derived cell lines expressing mutated P53 such as U373MG, U118MG or T98G cells (Ishii *et al*., 1999) failed to inhibit cell proliferation (Fig S1C) even though these cells do express plexin-A2 receptors (Fig. S1D). Mutations in p53 were found to enhance the progression of glioblastoma multiforme (Zhang *et al*., 2018) and it is thus likely that loss of plexin-A2 expression may not result in the inhibition of cell proliferation in glioblastoma cell lines that express mutated p53 or have lost p53 expression. Single pass receptors such as tyrosine-kinase receptors or plexins usually form homo or hetero dimers in order to transduce signals (Kigel *et al*., 2011;Sabag *et al*., 2014). In order to disrupt the dimerization dependent hypothetical pro-proliferative signaling of plexin-A2, we also expressed in the U87MG cells a cDNA encoding a dominant-negative truncated plexin-A2 lacking the intracellular domain of plexin-A2 (A2ExTm) (Fig. S2A). The expression of this cDNA also inhibited the proliferation of the cells significantly suggesting that homo or hetero dimerization of plexin-A2 may indeed be required for its pro-proliferative activity (Figs. S2B & S2C).

**Figure 1.**
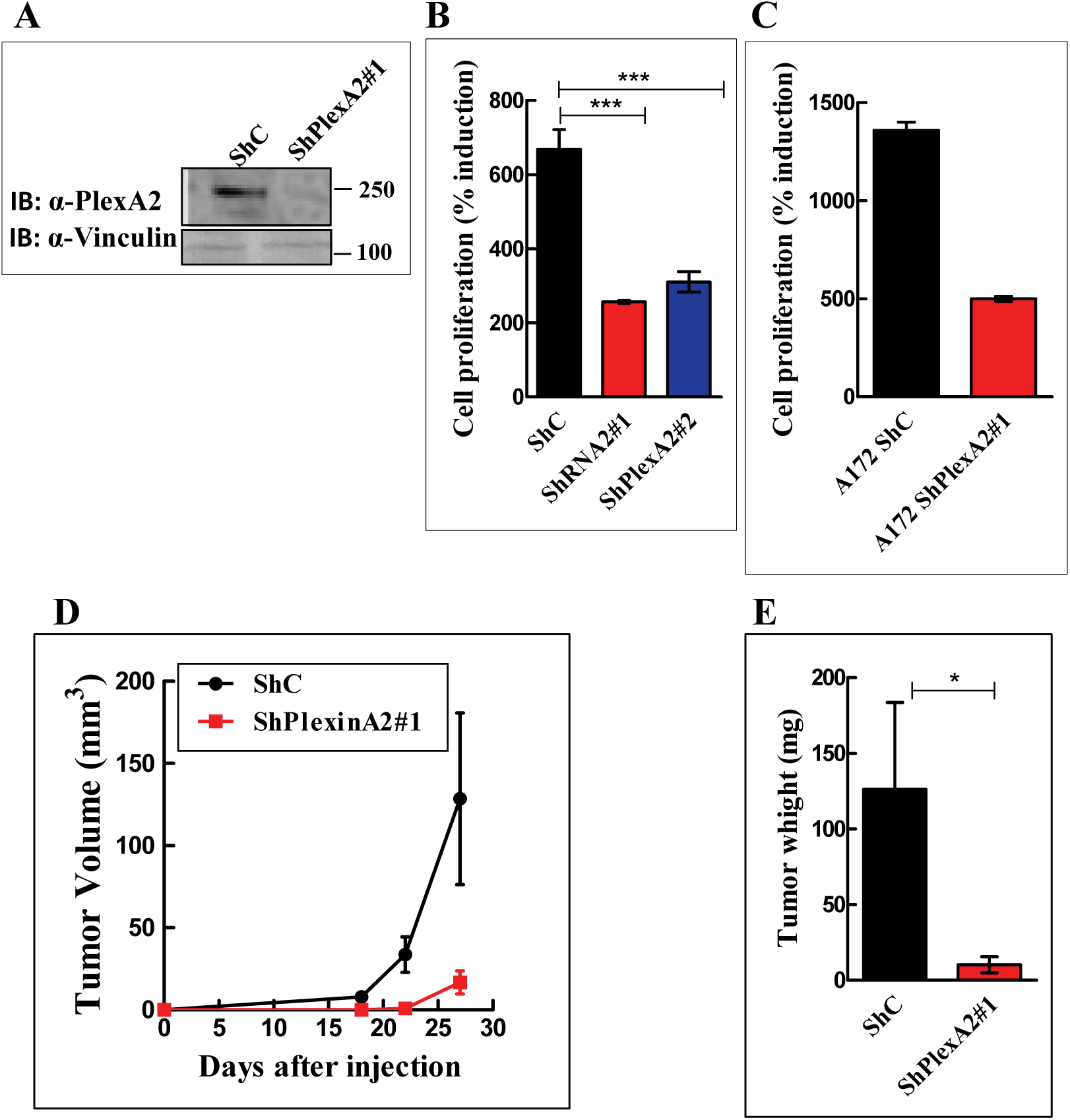
Silencing plexin-A2 expression in U87MG and A172 cells inhibits their proliferation and the development of subcutaneous tumors: **(A)** Western blot showing plexin-A2 expression levels in U87MG cells expressing a non-specific shRNA (ShC) or a shRNA targeting plexin-A2 (ShPlexA2#1). Cell lysates were probed with antibodies directed against plexin-A2 or vinculin. **(B)** U87MG or U87MG cells silenced for plexin-A2 expression using two different shRNAs were seeded in triplicate (2×10^4^ cells/well) in 24 well dishes. Cells were counted in a coulter counter after 3 days. Shown is the percentage of attached cells on day 3 compared with the number of cells attached on day 0 which was taken as 100%. Shown is the average of from three independent experiments. One-way ANOVA followed by Bonferroni’s multiple comparison post-test was used to assess statistical significance. Error bars represent the standard error of the mean. **(C)** A172 cells were infected with lentiviruses encoding a control shRNA (ShC) or a shRNA targeting plexinA2 (ShPlexA2#1). Each group of cells was seeded in quadruplicate in 96 well dishes (3×10^3^ cells/well). Cell proliferation was measured using the WST-1 proliferation assay as described in materials and methods, and the values presented were calculated as described under B. The experiment was repeated twice with similar results. Error bars represent the standard error of the mean. **(D)** U87MG expressing a non-specific shRNA (ShC) or cells that were silenced for plexin-A2 expression using ShPlexA2#1 were injected subcutaneously into Athymic/Nude mice (2×10^6^ cells/mouse). Tumor development was measured twice a week using calipers. **(E)** At the end of the experiment the tumors were excised and weighed. Each group contained 7 mice. The experiment was repeated twice with similar results. Statistical significance was evaluated using one tailed Mann-Whitney test. Error bars represent the standard error of the mean.

### Generation of U87MG cells lacking functional plexin-A2 receptor using CRISPR/Cas9

Silencing gene expression using shRNAs or the use of dominant negative inhibitors do not inhibit completely the expression of target genes. In order to determine the mechanisms by which plexin-A2 affects cell proliferation, we generated U87MG cells in which we have knocked-out the plexin-A2 gene by the introduction of frame-shift mutations into each of the alleles encoding plexin-A2 using CRISPR/Cas9. In order to differentiate as much as possible between true effects of plexin-A2 knock-out and off-target effects we used two different plexin-A2 specific guide RNAs (Fig. S3A) (Ran *et al*., 2013). We have subsequently isolated by limiting dilution several single cell derived clones of cells such as clones 35 and 54 that were generated using the different guide RNAs, in which each of the plexin-A2 encoding alleles contained frame shift mutations (Fig. S3B). These two clones no longer expressed plexin-A2 receptors (Fig. 2A) and have lost their ability to respond by cytoskeletal collapse to stimulation with sema3B (Fig. 2B) (Sabag *et al*., 2014). The proliferation of both knock-out clones was also very strongly inhibited as compared to the proliferation rate of U87MG cells (Fig. 2C-E) further suggesting that plexin-A2 is somehow able to promote the proliferation of U87MG cells.

**Figure 2.**
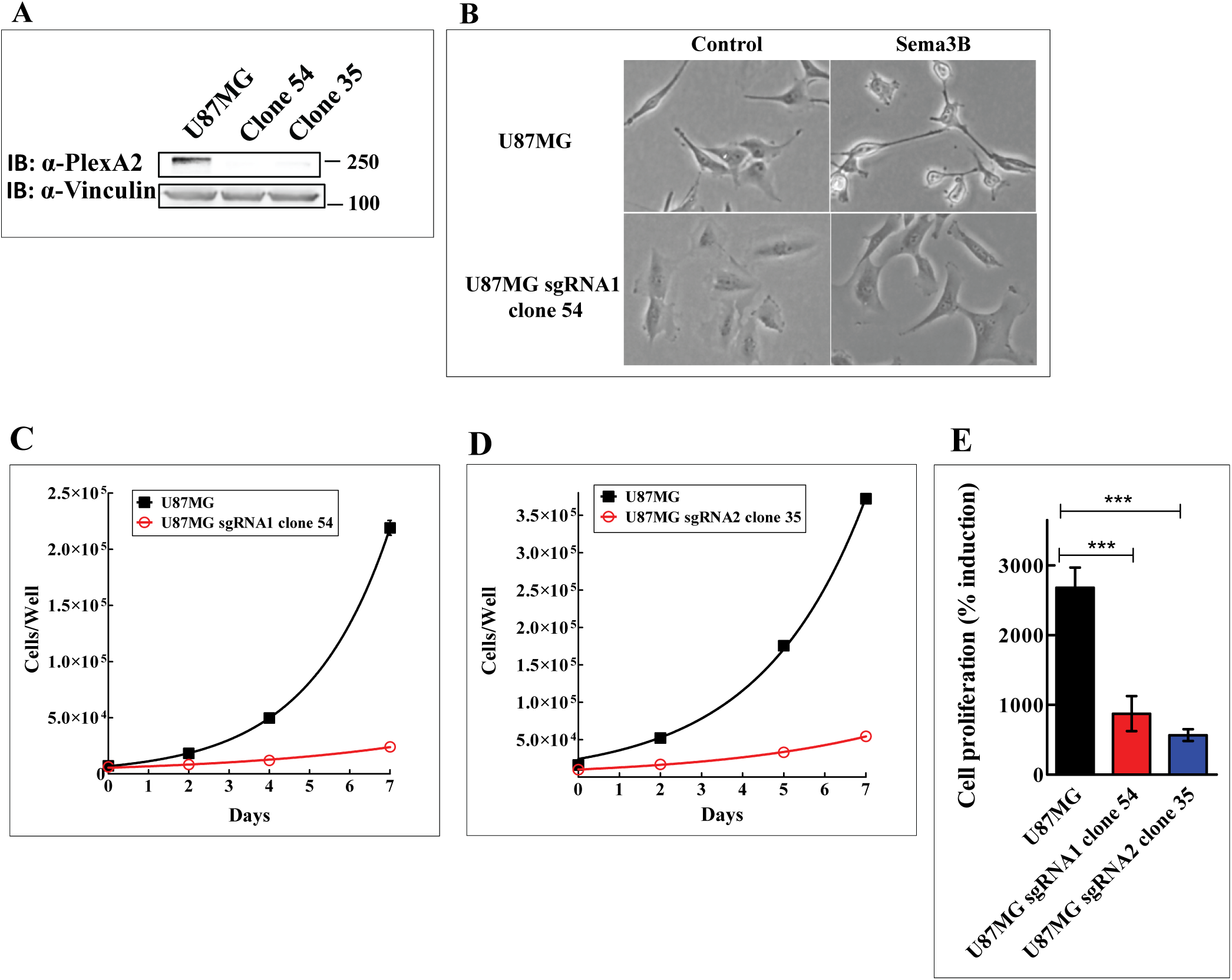
CRISPR/Cas9 mediated knock-out of plexin-A2 expression in U87MG cells inhibits their proliferation: **(A)** Two different guide RNAs were used to knock-out plexin-A2 expression in U87MG cells. Cell lysates of U87MG cells and knock-out cells (U87MG sgRNA1 clone 54 and U87MG sgRNA2 clone 35) were probed with antibodies directed against plexin-A2 or vinculin. **(B**) Conditioned medium from HEK293 cells infected with an empty lentiviral expression vector (Control) or from HEK293 cells expressing recombinant sema3B (300 μl) were added to parental U87MG cells or to U87MG sgRNA1 clone 54. Following a 30 min. incubation at 37°C the cells were photographed. **(C and D)** Representative growth curves of U87MG cells and plexin-A2 knock-out clone 54 and clone 35 cells. Cells were seeded in quadruplicate in complete growth medium (1×10^4^ cells/well). Aadherent cells were counted every two days using a coulter-counter. **(E)** The percentage of adherent cells on day 7 as compared to the number of cells attached on day 0 which was taken as 100%. Compared were parental U87MG cells (13 independent experiments), plexin-A2 clone 54 knock-out cells (8 independent experiments) and plexin-A2 clone 35 knock-out cells (5 independent experiments). One-way ANOVA followed by Bonferroni’s multiple comparison post-test was used to evaluate statistical significance. Error bars represent the standard error of the mean.

### The proliferation of plexin-A2 knock-out U87MG cells is rescued following re-expression of the plexin-A2 cDNA

In order to verify that the inhibition of cell proliferation observed following the knock-out of the plexin-A2 gene in U87MG cells or following shRNA mediated expression silencing of plexin-A2 is not due to off-target effects, we re-expressed the full-length cDNA encoding plexin-A2 in cells of the U87MG derived plexin-A2 knock-out clone 54. The proliferation of these cells was almost completely rescued following the re-expression of the plexin-A2 cDNA in three separate experiments (Fig. 3A). We also performed colony formation assays using parental U87MG cells, U87MG KO clone 54 cells and U87MG KO clone 54 cells in which we expressed the cDNA encoding the full length plexin-A2 cDNA. While clone 54 knock-out cells were almost completely unable to form colonies, re-expression of the plexin-A2 cDNA resulted in an almost complete rescue of colony formation ability (Fig. 3B).

**Figure 3.**
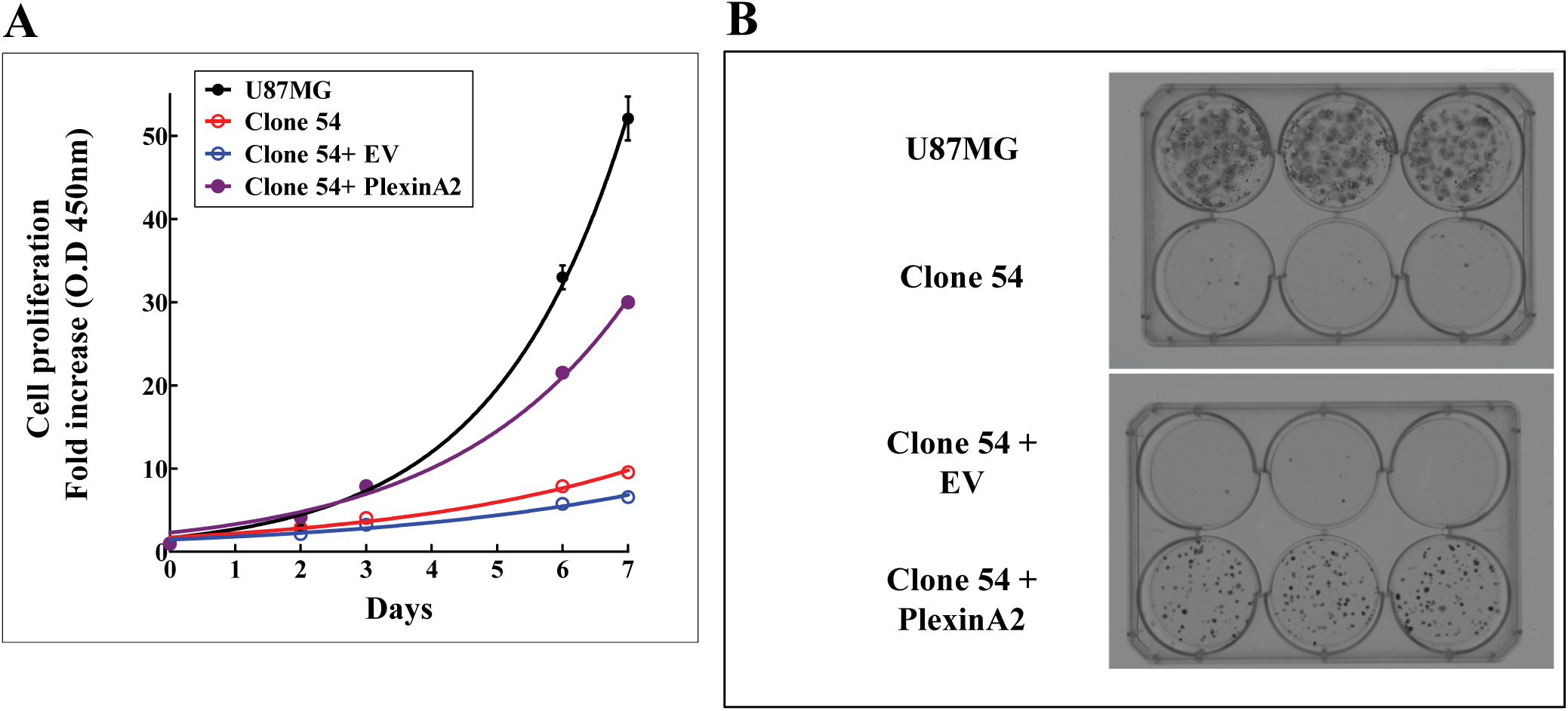
Re-expression of plexin-A2 cDNA in plexin-A2 knock-out cells, restores their proliferation and colony forming ability: **(A)** U87MG cells, U87MG clone 54 cell and clone 54 cells infected with empty lentiviruses (EV) or lentiviruses directing expression of plexin-A2 (PlexinA2) were seeded in quadruplicate in 96 well dishes. Cell proliferation was determined using the WST-1 assay as described. Shown is a representative experiment. **(B)** U87MG cells, U87MG clone 54 cells and U87MG KO clone 54 cells infected with an empty lentiviral expression vector (EV) or with a lentiviral vector directing expression of plexin-A2 (PlexinA2). Cells were seeded in 6 well plates in complete growth medium. After 16 days colonies were fixed in 4% para-formaldehyde and stained with crystal violet. The experiment was repeated three times with similar results.

### Expression of a truncated plexin-A2 lacking the extracellular domain restores cell proliferation of plexin-A2 knock-out cells

Clone 54 and clone 35 plexin-A2 knock-out cells proliferated at an extremely slow rate that made working with them very time consuming and difficult. We have therefore isolated, by limiting dilution, a subclone of the clone 54 cells which displayed a somewhat faster rate of basal proliferation (clone 54.3). Like clone 54 cells, the proliferation of clone 54.3 cells was also completely rescued following re-expression of the full length plexin-A2 cDNA (Fig. 4A-C). Expression of plexin-A2 in clone 54.3 cells, also enhanced the growth of anchorage independent colonies in soft agar which was strongly suppressed in the knock-out cells (Figs. 4D-F).

**Figure 4.**
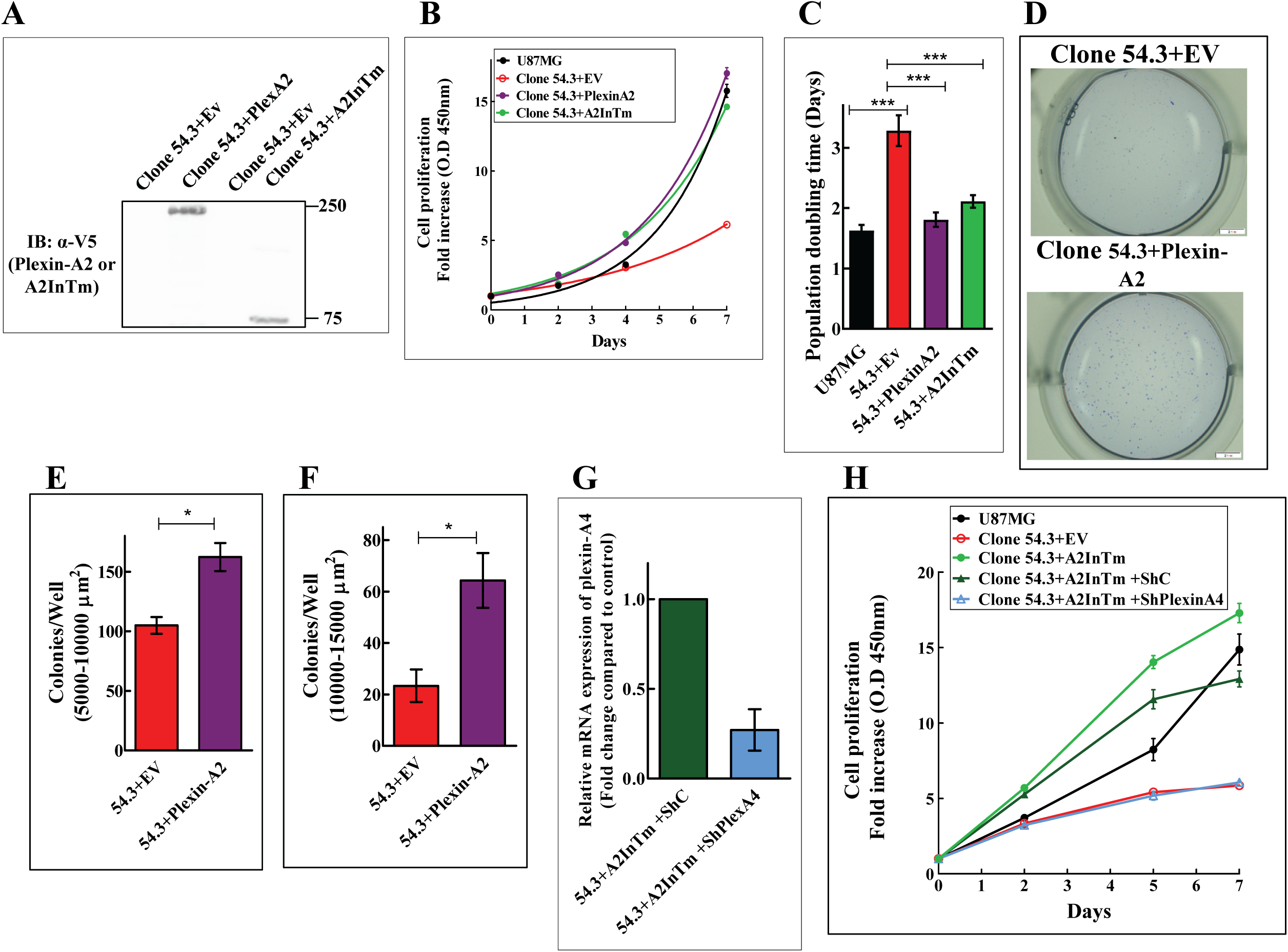
Re-expression of a truncated plexin-A2 containing the intracellular and trans-membrane domains (A2InTm) of plexin-A2 restores cell proliferation of plexin-A2 knock-out cells: **(A)** Plexin-A2 knock-out clone 54.3 cells were infected with an empty lentiviral expression vector (EV) or a lentiviral vector directing expression of a V5 tagged full length plexin-A2 (PlexA2) or a truncated plexin-A2 containing the intracellular and trans-membrane domains of plexin-A2 (A2InTm). Western blots prepared from cell lysates were probed with an antibody directed against V5. **(B)** Representative growth curves of U87MG cells and of U87MG KO clone 54.3 cells infected with lentiviruses encoding plexin-A2 (PlexinA2), A2InTm or an empty expression vector (EV). Cells were seeded in quadruplicate in 24 well dishes (1×10^4^ cells/well). Cells were counted every two days using a coulter-counter. **(C)** The average population doubling time of U87MG cells (N=11) was compared with the average population doubling time of clone 54.3+EV (N=11), clone 54.3+PlexinA2 (N=7) and clone 54.3+A2InTm (N=8). Statistical significance was evaluated using one-way ANOVA followed by Bonferroni’s multiple comparison post-test. Error bars represent the standard error of the mean. **(D)** Clone 54.3 cells infected with an empty lentiviral expression vector (EV) or with a lentiviral vector directing expression of plexin-A2 were seeded in soft agar in 24 well dishes (1×10^3^ cells/well) in triplicates. Colonies were stained with crystal violet after 13 days. Shown are photographs of representative wells. **(E and F)** The average number of colonies per well with an area between 5000-10000 µm^2^ or 10000-15000 µm^2^ were determined. Statistical significance was evaluated using the one tailed Mann-Whitney test. Error bars represent the standard error of the mean. **(G)** Lentiviruses were used to express control (ShC) or a plexin-A4 targeting shRNA (ShPlexA4) in KO clone 54.3 cells expressing A2InTm. The expression of plexin-A4 was then examined using qRT-PCR. Error bars represent the standard error of the mean of two independent experiments. **(H)** Representative growth curve of U87MG cells and of clone 54.3 cells expressing an empty expression vector (EV), A2InTm,. Clone 54.3 cells expressing A2InTm as well as a non-specific shRNA (ShC) or Clone 54.3 cells expressing A2InTm and a shRNA targeting plexin-A4 (ShPlexinA4). The various cells were seeded in quadruplicate in 96 well dishes in complete growth medium (3×10^3^ cells/well). The proliferation of the cells was determined every two days using the WST-1 assay as previously described. The experiment was repeated three times with similar results.

To identify plexin-A2 domains responsible for the transduction of its pro-proliferative signals in U87MG cells we first expressed in clone 54.3 cells a truncated plexin-A2 lacking the extracellular domain of plexin-A2 (A2InTm) (Fig. 4A). The expression of A2InTm also rescued efficiently the proliferation of the clone 54.3 cells (Figs. 4B & 4C), suggesting that elements in the intracellular domain are likely responsible for the transduction of the pro-proliferative activity of plexin-A2.

We have previously observed that the functional receptors for sema3A and sema3B are composed of a neuropilin, a plexin-A4 receptor, and either a plexin-A1 receptor in the case of sema3A or plexin-A2 in the case of sema3B (Kigel *et al*., 2011;Sabag *et al*., 2014). We therefore determined whether an additional type-A plexin may also be required for the pro-proliferative activity of plexin-A2 in U87MG cells. We have previously observed that silencing the expression of plexin-A4 in U87MG inhibits the proliferation of U87MG cells as well as their tumor forming ability (Kigel *et al*., 2011). It was recently observed that plexin-A2 and plexin-A4 form hetero dimers (Christie *et al*., 2021). Taken together these observations suggested that the pro-proliferative effects may be mediated by a plexin-A2/plexin-A4 receptor complex rather than by plexin-A2 alone. Indeed, we found that when we silence the expression of plexin-A4 in clone 54.3 plexin-A2 knock-out cells, expression of A2InTm alone is no longer sufficient to rescue the proliferation of the cells (Figs. 4G & 4H), indicating that both receptors are likely required for the pro-proliferative function.

### knocking-out plexin-A2 in U87MG cells results in cell cycle arrest and in the acquisition of some characteristics associated with senescent cells

Inhibition of cell proliferation as a result of the knock-out of the plexin-A2 gene in the U87MG cells was also accompanied by profound morphological changes. U87MG cells in which the plexin-A2 was knocked-out (clone 54) displayed a flattened morphology, and resembled in appearance and proliferation rate senescent cells (Fig 5A). Similar changes were also observed in clone 35 cells (Fig. 5A). U87MG cells in which we expressed the dominant-negative truncated plexin-A2 lacking the intracellular domain (A2ExTm) likewise have a flattened morphology (Fig. 5A). These morphological changes were completely reversed following the re-expression of the full length plexin-A2 cDNA or of the cDNA encoding A2InTm. The cells regained a polarized appearance, contracted, and appeared to be even smaller than parental U87MG cells, possibly because of plexin-A2 over-expression (Fig. 5A). In contrast with wild type U87MG cells, the knock-out cells stained positive for senescence associated beta-galactosidase (SA-β-gal) activity (Figs. 5B & 5D) (Dimri *et al*., 1995), suggesting that the knock-out cells display at least some characteristics of senescent cells, and further suggesting that plexin-A2 transduces in these cells signals that may inhibit cellular senescence. Similar morphological changes and staining for SA-β-gal activity were observed when the expression of plexin-A2 in U87MG cells or A172 glioblastoma cells was inhibited using shRNAs targeting plexin-A2 expression (Figs. 5C & S4). Surprisingly, the SA-β-gal activity was not suppressed following re-expression of full length plexin-A2 (Figs. 5B & 5D) or A2InTm (Fig. 5D) even though the proliferation of the cells had been rescued.

**Figure 5.**
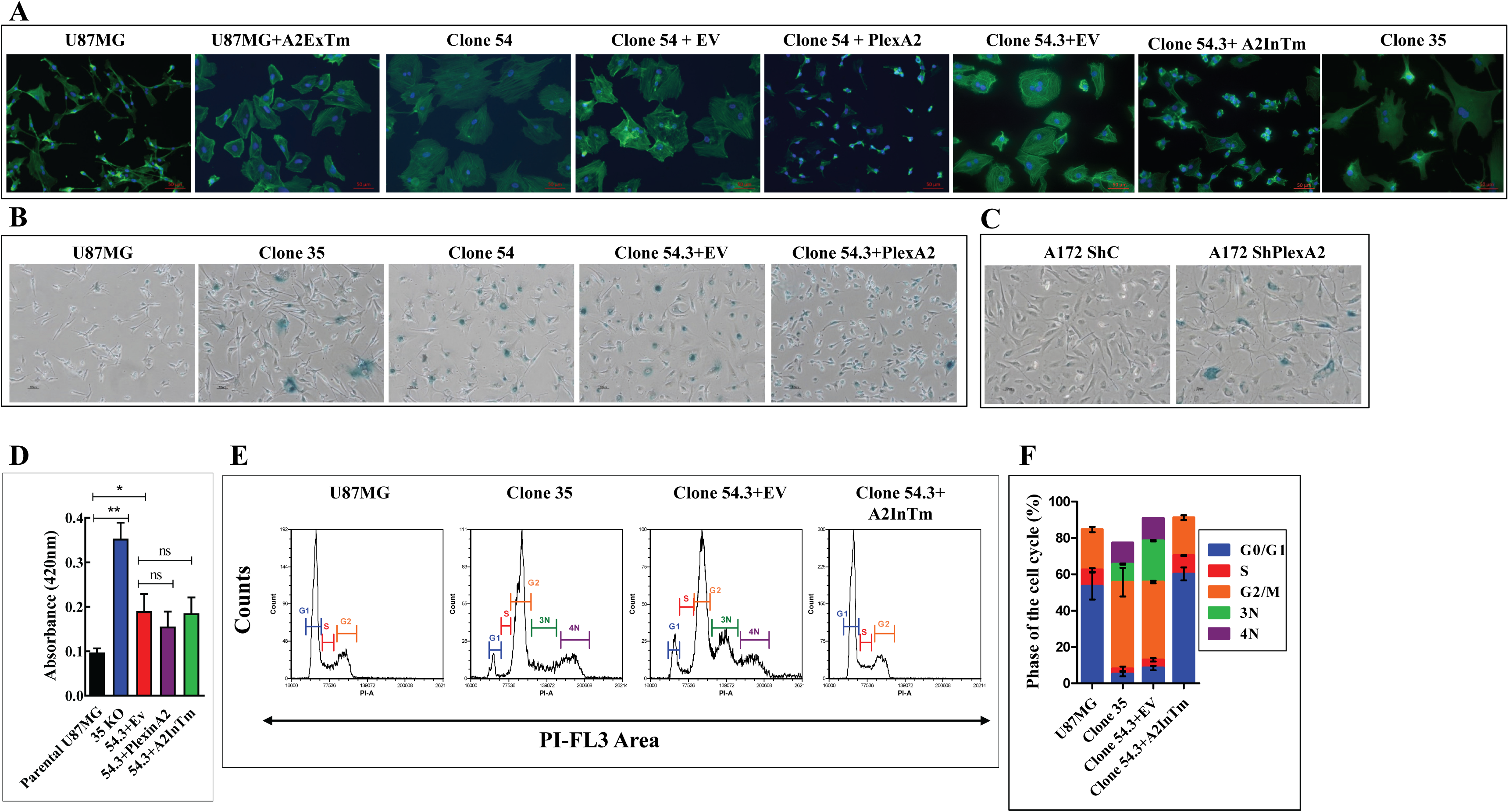
U87MG cells in which the plexin-A2 gene was knocked out acquire some properties of senescent cells: **(A)** The indicated cell types were seeded on fibronectin coated coverslips and stained with DAPI to visualize cell nuclei (blue) and with fluorescent phalloidin (green) to visualize actin fibers. Shown are merged confocal photographs generated using ZEN 2.3 software. **(B-C)** The indicated cell types were assayed at pH-6 for the expression of the senescence marker SA-β galactosidase as described in materials and methods. Shown are representative photographs **(D)** Equal numbers of U87MG cells (N=9 independent experiments), plexin-A2 knock out U87MG cells (clone 54.3 (N=7), clone 35 (N=3)) or clone 54.3 cells in which full length plexin-A2 (PlexinA2) (N=5) or A2InTm (N=5) were re-expressed, were collected and lysed in 100 µL of 0.1 M phosphate buffer (pH 6.0). β-galactosidase activity was then determined at pH 6.0 as described. Statistical significance was evaluated using the one tailed Mann-Whitney test. Error bars represent the standard error of the mean. **(E)** Cytofluorimetric analysis of propidium iodide-stained triplicates of U87MG cells, U87MG clone 35 cells and clone 54.3 cells infected with empty lentiviruses (EV) or A2InTm. The peaks represent cells in G0/G1 (G1), Cells in S (S), cells in G2/M (G2), and aneuploid cells (3N,4N). **(F)** The percentage of cells in the different cell cycle phases is shown for each of the cell types shown in E. Error bars represent the standard error of the mean (N=3).

Analysis of the cell cycle of clone 54.3 knock-out cells revealed a decrease in the fraction of cells found in the G0/G1 and S phases of the cell cycle which was accompanied by an increase in the percentage of cells found in the G2 phase as compared with wild type U87MG cells or with plexin-A2 knock-out cells in which we have expressed A2InTm (Figs. 5E & 5F). Furthermore, we could detect an increase in the percentage of aneuploid cells in plexin-A2 knock-out cells (Fig. 5E & 5F). These observations suggest that inhibition of plexin-A2 expression inhibits cell cycle progression in the G2/M phase.

### Inhibition of plexin-A2 expression is accompanied by decreased AKT phosphorylation and by enhanced phosphorylation of p38-MAPK

In order to identify secondary messengers that transduce of the pro-proliferative signals of plexin-A2 we identified known secondary messengers whose phosphorylation state changes following plexin-A2 knock-out in U87MG cells. These experiments revealed that the phosphorylation of AKT on ser473 is inhibited in plexin-A2 clone 54.3 knock-out cells (Figs. 6A & 6B) as well as in clone 35 knock-out cells (Figs. S5A & S5B). In contrast the phosphorylation of p38-MAPK was induced on Tyr182 in these cells (Figs. 6C, 6D, S5C and S5D). Enhanced p38-MAPK phosphorylation on Tyr182 was reported to be associated with Ras induced premature senescence (Wang *et al*., 2002;Han and Sun, 2007). The changes we observed in the phosphorylation states of p38 and AKT are specific to plexin-A2 since AKT phosphorylation is restored following expression of A2InTm in clone 54.3 knock-out cells while the phosphorylation of p38 is reduced to basal levels (Figs. 6A-6D). The restoration is not due to an artefact caused by infection stress since expression of an empty vector (Figs. 6A-6D) or of the dominant negative form of plexin-A2 which lacks the intracellular domain (Figs. 6A & 6B) did not reverse the phosphorylation state to pre-knock-out levels.

**Figure 6.**
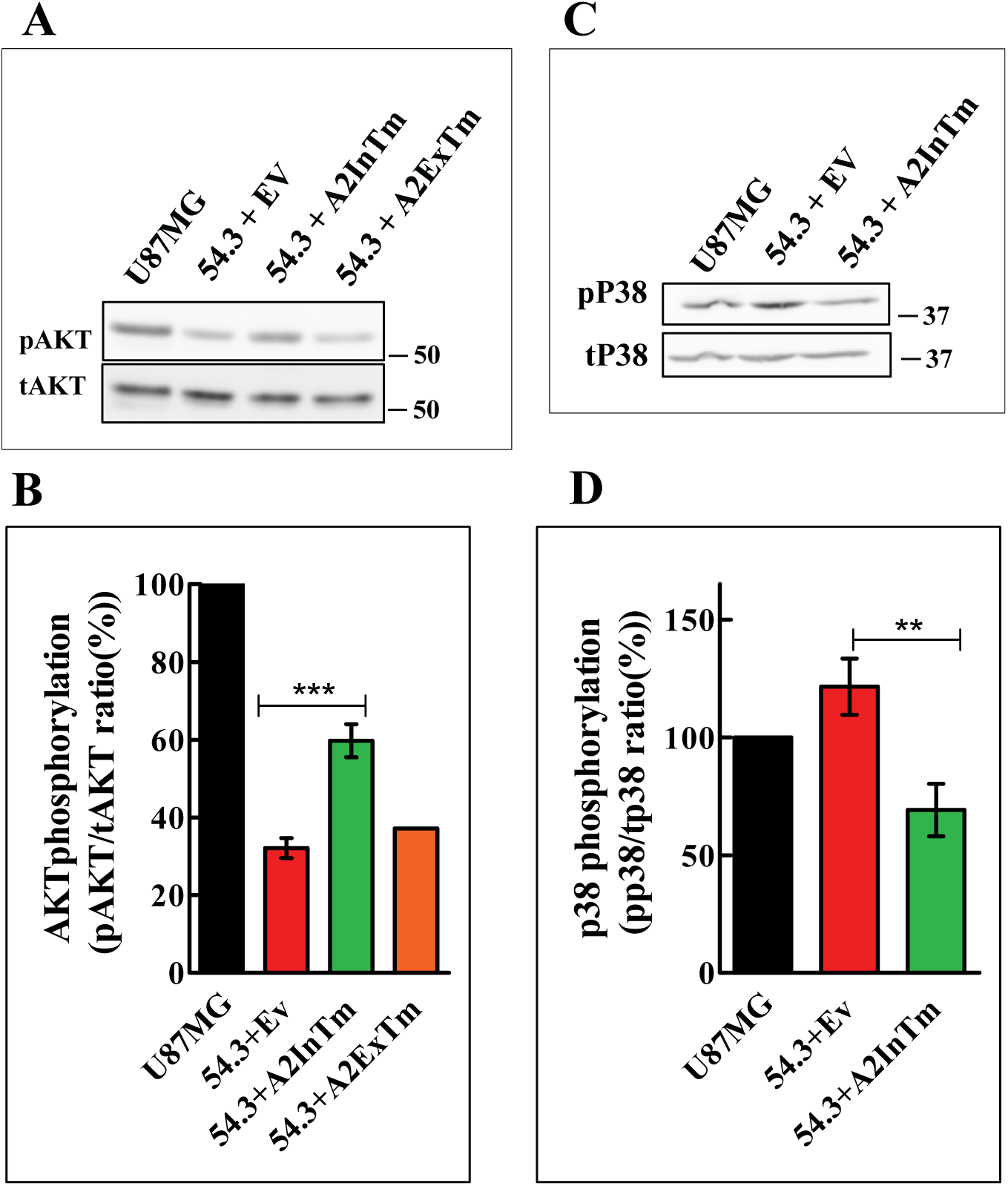
The phosphorylation of AKT is inhibited and that of p38 induced in U87MG cells in which the plexin-A2 gene is knocked-out: The cDNAs encoding A2InTm, A2ExTM or an empty expression vector (EV) were expressed in clone 54.3 knock-out cells (54.3). **(A)** The phosphorylation levels of AKT were assayed using western blot analysis of cell lysates and an antibody directed against phosphorylated AKT (ser473). Loading was assessed using an antibody directed against total AKT. Shown is a representative western blot out of 8 similar experiments **(B)** The effect of A2InTm expression on the average phosphorylation level of AKT was determined in 8 independent experiments (except for A2ExTm expressing cells, N=1) as described. Shown is the average phosphorylation level of AKT. Statistical significance was evaluated using one-way ANOVA followed by Bonferroni’s multiple comparison post-test. Error bars represent the standard error of the mean. **(C)** P38 phosphorylation levels were assayed in the indicated cell types by western blot analysis of cell lysates using an antibody directed against phosphorylated p38 (Thr180/Tyr182). Loading was assessed using an antibody directed against total p38. Shown is a representative experiment. **(D)** The effect of A2InTm expression on the average phosphorylation level of p38 derived from 6 independent experiments is shown. Statistical significance was evaluated using one-way ANOVA followed by Bonferroni’s multiple comparison post-test. Error bars represent the standard error of the mean.

### The FARP2 binding site and the FYN dependent phosphorylation sites of plexin-A2 mediate the pro-proliferative activity of plexin-A2

To identify plexin-A2 intracellular sub-domains that mediate the pro-survival/pro-proliferative activity of plexin-A2, we introduced point mutations into four known functional sub-domains in the plexin-A2 intracellular domain. These included the Rho GTPase binding domain (RBD domain) (Wang *et al*., 2011;Hota *et al*., 2012;Zhang and Buck, 2017), the KRK motif of the FERM domain that participates in the binding of FARP2 (Toyofuku *et al*., 2005;Mlechkovich *et al*., 2014), the tyrosine residues that are phosphorylated by the FYN tyrosine-kinase (St Clair *et al*., 2017) and the juxtamembrane cytosolic dimerization interface domain) that was found to contribute to plexin-A3 homo-dimerization (Barton *et al*., 2015) (Fig. S6A).

The cDNAs encoding the mutated A2InTm variants were expressed in clone 54.3 knock-out cells (Fig. S6B) and assayed for their ability to rescue cell proliferation. The mutations that were introduced into the RBD domain and into the juxtamembrane cytosolic dimerization interface did not inhibit the rescuing ability of A2InTm (Figs. 7A). In contrast, the mutations introduced into the FARP2 binding site and into the FYN phosphorylation sites each partially inhibited the rescue by A2InTm (Figs. 7A & 7B). The introduction of mutations into the FARP2 binding site also inhibited strongly A2InTm’s ability to rescue the formation of anchorage independent colonies from single cells in soft agar. In contrast, the formation of colonies in soft agar from cells expressing A2InTm containing the FYN phosphorylation sites mutations was not inhibited significantly although a small partial effect that did not reach statistical significance could still be observed (Fig. 7C-E).

**Figure 7.**
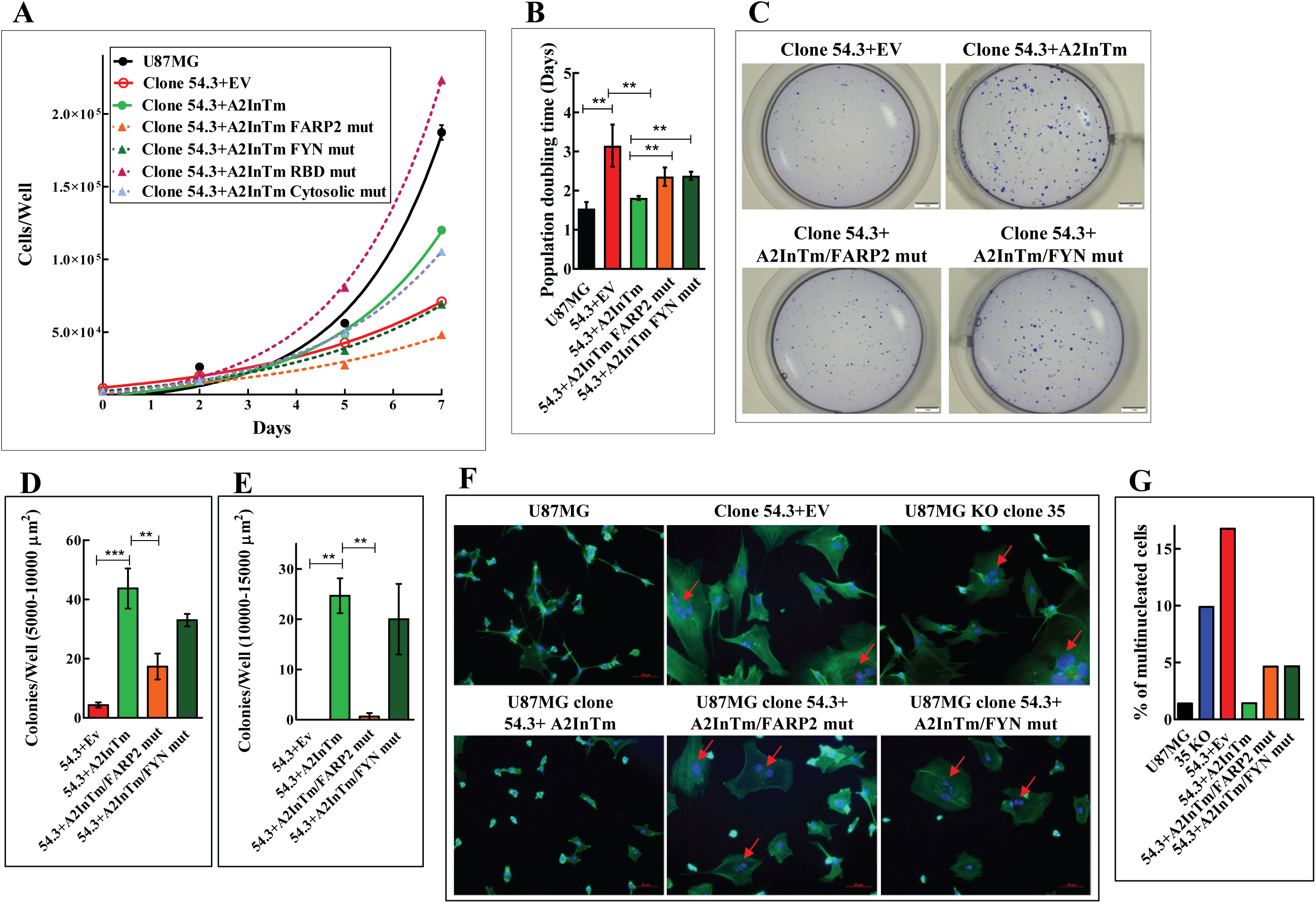
Point mutations in the FARP2 binding site and in the Fyn phosphorylation sites of the plexin-A2 intracellular domain inhibit restoration of cell proliferation induced by A2InTm: **(A)** U87MG cells, U87MG clone 54.3 cells infected with either control lentiviruses (EV) or with lentiviruses encoding A2InTm or A2InTm variants containing point mutations in the FARP2 binding site (54.3+A2InTm FARP2 mut), in the Fyn phosphorylation sites (54.3+A2InTm FYN mut), in the Rho binding domain (RBD) (54.3+A2InTm RBD mut) or in the juxtamembrane cytosolic dimerization interface (54.3+A2InTm Cytosolic mut) of plexin-A2 were seeded in 24 well dishes (1×10^4^ cells/well). Triplicate wells were counted every two days using a coulter-counter. Shown is a representative experiment out of five similar experiments. **(B)** Shown are the average population doubling times of the clone 54.3 derived indicated cell types. Results were derived from 5 independent experiments. Statistical significance was evaluated using the one tailed Mann-Whitney test. Error bars represent the standard error of the mean. **(C)** Single cell suspensions of clone 54.3+EV, clone 54.3+A2InTm, clone 54.3+A2InTm FARP2 mut or clone 54.3+A2InTm FYN mut cells, were seeded in soft agar (1×10^3^ cells/well) in triplicates. After 17 days colonies of cells were stained with crystal violet. Shown are photographs of representative wells. **(D and E)** The average number of colonies per well with an area between 5000-10000 µm^2^ or 10000-15000 µm^2^ was then determined. Statistical significance was evaluated using one-way ANOVA followed by Bonferroni’s multiple comparison post-test. Error bars represent the standard error of the mean. **(F)** U87MG cells, clone 54.3+EV cells, clone 35 cells, clone 54.3+A2InTm cells, clone 54.3+A2InTm FARP2 mut cells and clone 54.3+A2InTm FYN mut cells were seeded on fibronectin coated cover slips and stained with DAPI to visualize cell nuclei (blue) and with fluorescent phalloidin (green) to visualize actin fibers. Shown are merged confocal photographs generated using ZEN 2.3 software. Red arrows point at multinucleated cells. **(G)** The percentage of multinucleated cells of the various cell types described in F was determined by counting 23 microscopic fields per each cell type.

A2InTm containing mutations in the FARP2 binding site as well as A2InTm containing mutations in the FYN phosphorylation sites also lost the ability to reverse the morphological changes observed in plexin-A2 knock-out cells (Fig. 7F). We have also found that plexin-A2 knock-out cells contain a higher percentage of multinucleated cells. The concentration of multinucleated cells was strongly reduced following the expression of A2InTm in the knock-out cells. Expression of A2InTm mutated in the FARP2 binding site or in the FYN phosphorylation sites each resulted in a partial decrease in the percentage of multinucleated cells, which was less efficient than the rescue obtained with wild type A2InTm (Figs. 7F & 7G) suggesting that the function of both sites is probably required for complete rescue.

### The FARP2 binding site and the FYN phosphorylation sites of plexin-A2 are required for the phosphorylation of AKT

The phosphorylation of AKT on ser473 is inhibited in plexin-A2 knock-out U87MG cells and rescued by A2InTm expression (Figs. 6A & 6B). Interestingly, mutations in the FARP2 binding site or in the FYN phosphorylation sites also inhibited partially the rescue of AKT phosphorylation suggesting that AKT may indeed mediate, at least in part, the transduction of the pro-proliferative effects of plexin-A2 (Figs. 8A, 8B, 8C and 8D). Indeed, FYN was previously reported to activate AKT to promote glioma cell proliferation (Law and Lee, 2012).

**Figure 8.**
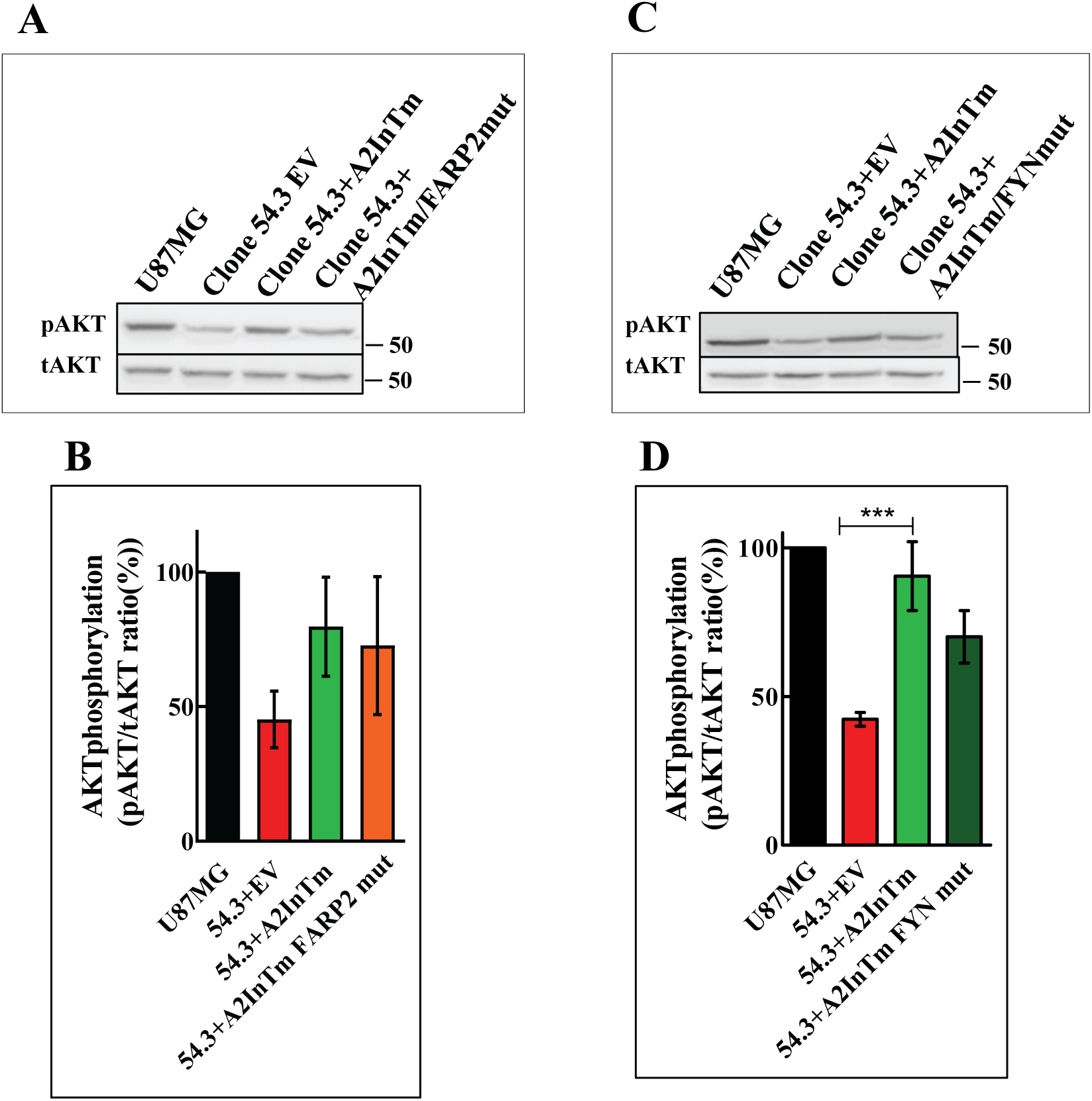
Point mutations in the FARP2 binding site and the Fyn phosphorylation sites of plexin-A2 inhibit the A2InTm mediated rescue of AKT phosphorylation: An empty letiviral expression vector (EV) or lentiviruses expressing cDNAs encoding A2InTm or A2InTm variants containing point mutations in the FARP2 binding site (A2InTm FARP2 mut) or in the Fyn phosphorylation sites (A2InTm FYN mut)) were expressed in clone 54.3 knock-out cells (54.3). **(A)** The cells were lysed and the levels of AKT phosphorylation determined by western blot analysis as described in methods. Shown is a representative experiment. **(B)** The effect of expression of A2InTm variant containing point mutations in the FARP2 binding site (A2InTm FARP2 mut) on the average phosphorylation level of AKT was determined in 5 independent experiments as described in methods. Statistical significance was evaluated using one-way ANOVA followed by Bonferroni’s multiple comparison post-test. Error bars represent the standard error of the mean. **(C)** The indicated cell types were lysed and the levels of AKT phosphorylation were determined in western blots as described in the methods. Shown is a representative experiment. **(D)** The effect of expression of A2InTm variant containing point mutations in the Fyn phosphorylation sites (A2InTm FYN mut) on the average phosphorylation level of AKT derived from 6 independent experiments is shown. Statistical significance was evaluated using one-way ANOVA followed by Bonferroni’s multiple comparison post-test. Error bars represent the standard error of the mean.

### Plexin-A2 mediated proliferation of U87MG glioblastoma cells is independent of sema3C

It was reported that inhibition of sema3C expression in glioblastoma cancer stem cells inhibits cell proliferation and it was suggested that this activity is mediated by the plexin-D1 or the plexin-A2 receptors (Man *et al*., 2014). However, we could not detect any sema3C in the conditioned medium of U87MG cells (Fig. 9A). Sema3E signals exclusively using the plexin-D1 receptor (Gu *et al*., 2005). U87MG cells do not express plexin-D1 receptors (Fig. S1D) (Smolkin *et al*., 2018), and did not respond to stimulation with sema3E by cell contraction (Fig. 9B) (Casazza *et al*., 2012;Smolkin *et al*., 2018). Activation of plexin-A2 mediated signal transduction by sema3C would require the binding of sema3C to the neuropilin-1 or neuropilin-2 receptors which then associate with plexin-A2 to initiate signal transduction (Rohm *et al*., 2000;Janssen *et al*., 2012;Hao and Yu, 2018;Smolkin *et al*., 2018;Toledano *et al*., 2019). It follows that if the knock-out of plexin-A2 inhibits the proliferation of U87MG cells due to the disruption of plexin-A2 mediated sema3C autocrine signal transduction, then it is expected that inhibition of the expression of the neuropilin receptors should also result in inhibition of cell proliferation. However, the proliferation of U87MG cells in which we have knocked-out the genes encoding neuropilin-1 and neuropilin-2 using CRISPR/Cas9 (Smolkin *et al*., 2018) is not inhibited (Figs. 9C & 9D) suggesting that the inhibition of cell proliferation induced by the knock-out of plexin-A2 is not due to the disruption of sema3C induced autocrine signaling.

**Figure 9.**
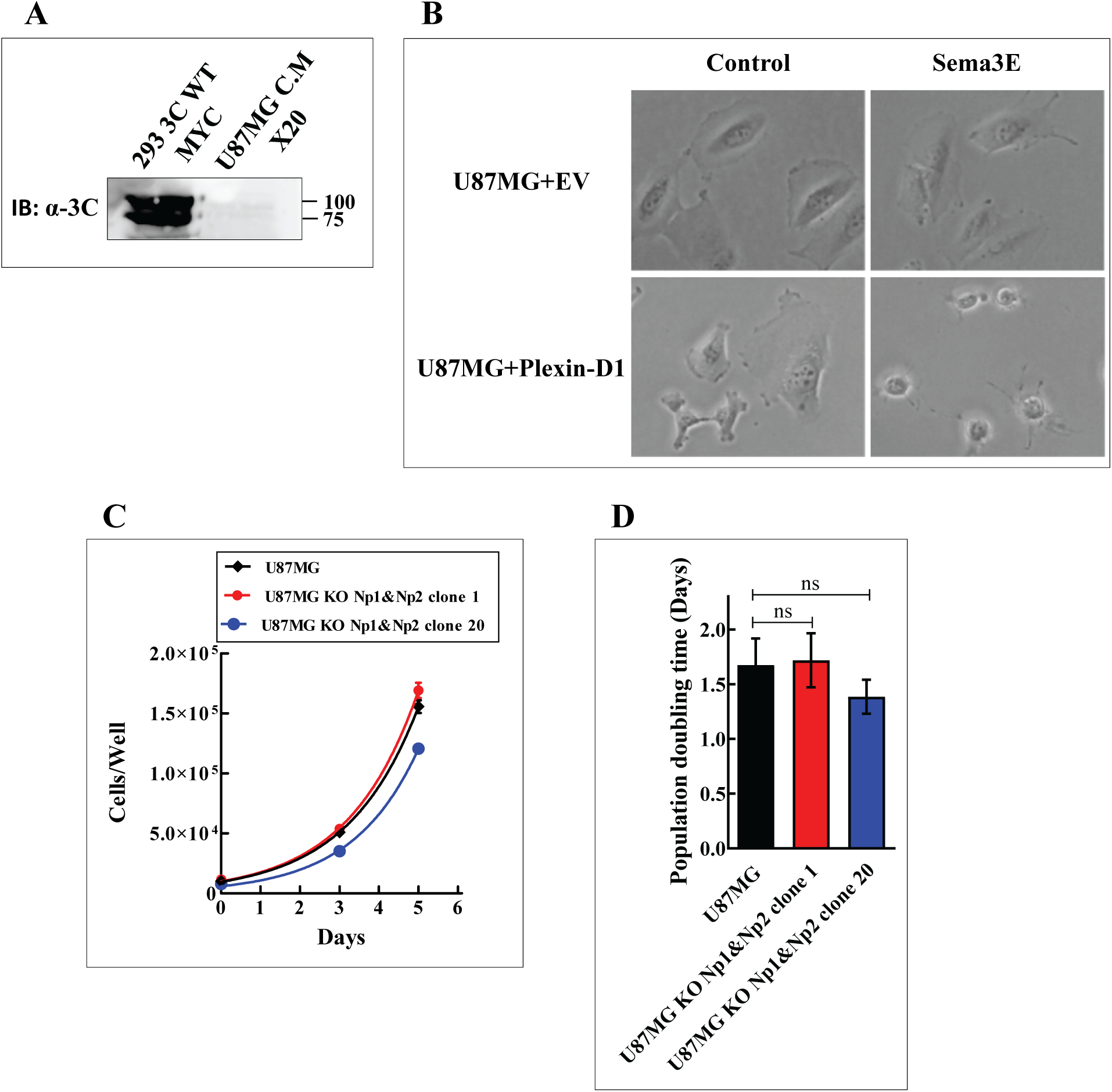
The pro-proliferative effects of plexin-A2 are not induced by sema3C: **(A)** Conditioned medium of U87MG cells was concentrated 20-fold by centrifugation using Amicon™ 30K centrifuge filters. Shown is a western blot probed with an antibody directed against sema3C. A cell lysate prepared from HEK293 cells over-expressing sema3C (Mumblat *et al*., 2015) was used as a positive control. **(B)** U87MG cells infected with an empty lentiviral expression vector (U87MG+EV) as well as U87MG cells expressing recombinant plexin-D1 (U87MG+Plexin-D1) (Smolkin *et al*., 2018), were stimulated with elution buffer (Control) or with 1 µg/ml purified sema3E/Fc (Casazza *et al*., 2012). Following a 30 min. incubation at 37°C the cells were photographed. **(C)** Representative growth curves of U87MG cells and of two independent U87MG derived clones of cells in which both the NRP1 and NRP2 genes were knocked-out (Smolkin *et al*., 2018). Cells were seeded in complete growth medium (1×10^4^ cells/well). Aadherent cells were counted every two days using a coulter-counter. **(D)** The average population doubling times of the cell types shown in C derived from five independent experiments were then evaluated for statistical significance using one-way ANOVA followed by Bonferroni’s multiple comparison post-test. Error bars represent the standard error of the mean.

## DISCUSSION

Plexins such as plexin-B1, plexin-B2 and plexin-A4 were recently found to function as transducers of pro-tumorigenic and pro-proliferative signals conveyed by semaphorins such as, sema4D, sema4C and sema6B (Conrotto *et al*., 2005;Kigel *et al*., 2011;Gurrapu *et al*., 2018). Plexin-A2 was previously reported to convey anti-angiogenic signals induced by sema3B (Varshavsky *et al*., 2008) and to be down regulated in tumors from several types of cancer (Gabrovska *et al*., 2011;Valiulyte *et al*., 2020). In another report plexin-A2 was implicated as a mediator of glioblastoma tumor progression induced by sema3C (Man *et al*., 2014) but a recent report of the same group reversed that conclusion (Christie *et al*., 2021). Here we have found that silencing the expression of the plexin-A2 receptor in U87MG and A172 glioblastoma cells as well as in human umbilical vein derived endothelial cells (HUVEC) using specific shRNA species resulted in very strong inhibition of cell proliferation. Furthermore, silencing the expression of the plexin-A2 receptor in U87MG glioblastoma cells resulted also in very strong inhibition of tumor development from these cells. Similar results were obtained when we inhibited plexin-A2 function using a truncated dominant negative form of plexin-A2 and when we knocked-out the plexin-A2 gene in these cells using CRISPR/Cas9. In contrast with the U87MG and A172 cells, silencing the expression of plexin-A2 in U373MG, U118MG or T98G glioblastoma multiforme derived cells did not affect their proliferation rate. Interestingly, in contrast with U87MG and A172 cells, these three cell lines contain mutated p53 (Ishii *et al*., 1999) suggesting that plexin-A2 may be somehow involved in pathways that regulate the activity of P53. This will need to be further examined in the future.

Silencing gene expression by shRNAs or dominant negative constructs does not usually inhibit gene expression completely while gene knock-out does. In order to gain a better understanding of the molecular mechanisms by which plexin-A2 promotes cell proliferation, we therefore used U87MG single cell derived clones of cells in which we have knocked-out the plexin-A2 gene in both alleles by the introduction of frame shift mutations in the second exon of plexin-A2. The proliferation of these knock-out cells was extremely slow, and the clones tended to stop proliferating completely after a while. This was indeed due to inhibition of plexin-A2 expression since rescue experiments in which we re-expressed the full-length cDNA encoding plexin-A2 in the knock-out cells fully restored their proliferation. The loss and rescue of plexin-A2 expression in these cells was also accompanied by cell shape changes. The plexin-A2 knock-out cells as well as U87MG cells expressing truncated dominant negative form of plexin-A2 or an shRNA targeting the plexin-A2 mRNA adopted a flattened morphology reminiscent of the shape changes associated with senescent cells while re-expression of plexin-A2 restored their polarized morphology. Indeed, in contrast with parental U87MG cells, the knock-out cells also expressed senescence-associated beta-galactosidase (SA-β-gal) activity (Dimri *et al*., 1995). Similarly, Plexin-A2 silenced A172 glioblastoma cells also displayed SA-β-gal activity. Unexpectedly, the SA-β-gal activity persisted in rescued cells in which plexin-A2 was re-expressed even though the proliferation of the rescued cells was restored, and they certainly did not resemble senescent cells anymore. However, similar observations had been reported in the past (Piechota *et al*., 2016;Bojko *et al*., 2019).

Sema6A, sema6B and sema3B transduce their signals using plexin-A2 and plexin-A4 (Suto *et al*., 2007;Sabag *et al*., 2014). When the expression of sema6B is inhibited in endothelial cells or in U87MG the cells flatten, and their proliferation is inhibited in a manner reminiscent of the changes observed when plexin-A2 expression is inhibited in U87MG cells (Kigel *et al*., 2011). Since rescue experiments using the original plexin-A2 knock-out clones were very difficult due to their extremely slow proliferation rate, we isolated a partial spontaneous revertant U87MG knock-out clone (clone 54.3) which proliferated at a somewhat faster rate making re-expression of various construct a bit easier technically. The proliferation of clone 54.3 cells could still be rescued completely following the expression of the plexin-A2 cDNA. Using these cells, we found that the proliferation as well as the morphological changes can also be restored to wild type levels following the expression of the plexin-A2 deletion mutant A2InTm which lacks the entire extracellular domain. Furthermore, while the knock-out cells contained an elevated concentration of aneuploid cells no such cells could be detected in rescued cells.

It is so far unclear if plexin-A2 signaling actually promotes cell proliferation or only supplies a permissive signal that allows cell proliferation and protects against senescence. These questions will need to be further investigated in the future. However, our experiments clearly suggested that domains in the intracellular part of plexin-A2 are probably responsible for the pro-proliferative function of plexin-A2 in these cells. Indeed, we have found that point mutations introduced into the FARP2 binding domain of plexin-A2 (Toyofuku *et al*., 2005;Mlechkovich *et al*., 2014) inhibited partially rescue of cell proliferation as well as the reversal of the morphological changes by A2InTm expression. Similarly point mutations introduced into the two FYN phosphorylation sites of the intracellular domain (St Clair *et al*., 2017) also inhibited partially rescue of cell proliferation and inhibited as well as the reversal of the morphological changes by A2InTm, suggesting that both of these sites are probably required for the full pro-proliferative function of plexin-A2. Expression of plexin-A2 or of A2InTm in knock-out cells also restored the growth of anchorage independent colonies in soft agar which was almost completely abolished in clone 54.3 plexin-A2 knock-out cells. While Mutations in the FYN phosphorylation sites of the intracellular domain only had a marginal effect the ability to form colonies in soft agar, mutating the FARP2 domain abolished much more effectively rescue by A2InTm expression suggesting that FARP2 is much more important for anchorage independent proliferation which is one of the cancer cell hallmarks.

In order to identify potential secondary messengers that convey the pro-proliferative signal of plexin-A2 we screened for secondary messengers whose phosphorylation state is altered in the knock-out cells. We found that the phosphorylation of p38-MAPK was enhanced on tyr182 in the plexin-A2 knock-out cells while the phosphorylation of AKT on ser473 was inhibited. The phosphorylation of AKT was rescued following A2InTm expression in the cells but failed partially when the rescue was done using A2InTm containing mutations in the FARP2 binding domain or in the FYN phosphorylation sites of plexin-A2. Similarly, the phosphorylation state of p38MAPK was also rescued following A2InTm expression but in this case the rescue was not affected when A2InTm containing mutations in the FARP2 site or in the FYN phosphorylation sites was used to rescue the p38MAPK phosphorylation state, suggesting that p38MAPK is not linked to plexin-A2 via FYN or FAERP2. Increased phosphorylation of p38MAPK was previously reported to be associated with Ras induced premature senescence (Wang *et al*., 2002) and inhibition of AKT was also previously linked to inhibition of glioblastoma progression. (Bao *et al*., 2019). It was also previously observed that during selective apoptosis in glioblastoma cell lines caused by the MG132 proteasome inhibitor AKT activity is inhibited and p38 signaling is activated (Zanotto-Filho *et al*., 2012). It is thus possible that these two secondary messengers convey at least part of the plexin-A2 induced pro-proliferative signal transduction.

The pro-proliferative/pro-survival function of plexin-A2 may be independent of external activation or be activated by an autocrine factor that is constitutively expressed in these cells. It is possible that sema3C is somehow associated with these observations since it was reported that inhibition of sema3C in glioblastoma cancer stem cells also inhibits cell proliferation and that this is mediated by the plexin-D1 and by the plexin-A2 receptors (Man *et al*., 2014). However, U87MG cells do not express the sema3C plexin-D1 receptor and therefore fail to respond to stimulation with sema3E which signals exclusively using plexin-D1 receptors (Gu *et al*., 2005;Sabag *et al*., 2014). Even overexpression of plexin-A2 in U87MG cells does not enable signal transduction of sema3C in the absence of plexin-D1 (Smolkin *et al*., 2018). To enable the activation of plexin-A2 mediated signal transduction by class-3 semaphorins, they have to bind first to one of the neuropilin receptors (Neufeld *et al*., 2016). However, U87MG glioblastoma cells in which we knocked-out the genes encoding both neuropilins using CRISPR/Cas9 did not respond to sema3C, either by cytoskeletal collapse or by inhibition of proliferation. Thus it seems that the pro-proliferative activity of plexin-A2 is not the result of stimulation by autocrine sema3C. It is possible that other semaphorins or some other factor produced in an autocrine manner may be responsible for the pro-proliferative function of plexin-A2. We have found previously that plexin-A4 silencing also inhibits the proliferation of U87MG cells and tumor formation from these cells (Kigel *et al*., 2011). Indeed, when the expression of plexin-A4 was silenced in plexin-A2 knock-out cells, A2InTm expression failed to rescue cell proliferation. It therefore seems that the transduction of the pro-proliferative signal of plexin-A2 also requires plexin-A4. Indeed, plexin-A2 was found to be associated spontaneously physically with plexin-A4 in a recent publication (Christie *et al*., 2021). We have also previously found that silencing the expression of the plexin-A2/plexin-A4 ligand sema6B in U87MG cells also results in the inhibition of proliferation and tumor formation (Kigel *et al*., 2011). It is therefore likely that autocrine sema6B activates a pro-proliferative/pro-survival signal that is mediated by a hetero-dimeric complex of plexin-A2 and plexin-A4, and by secondary mediators such as FARP2, FYN, AKT and p38MAPK. Our results suggest a novel role for plexin-A2 as a modulator of glioblastoma cells proliferation and indicate that inhibitors of plexin-A2/plexin-A4 may perhaps represent novel targets for the development of anti-tumorigenic drugs.

## MATERIALS AND METHODS

### Antibodies and recombinant proteins

Mouse sema3E/FC was produced and purified as previously described (Casazza *et al*., 2012). Rabbit anti plexin-A2: Abcam (ab39357), Rabbit anti plexin-A4: Sigma (HPA029919), Goat anti plexin-D1: Abcam (ab28762), Mouse anti-vinculin: Chemicon (3574), Goat anti plexin-A2-APC: RαD systems (FAB5486A), Alexa Fluor 488 Phalloidin: Molecular Probes (A12379), Mouse anti-V5: Invitrogen (R960-25), Rabbit anti phospho-AKT (ser473): Cell Signaling (4060), Mouse anti AKT: Santa-Cruz (sc-5298), Rabbit anti phospho-p38 MAPK (Thr180/Tyr182): Cell Signaling (#9211), Rabbit anti p38-MAPK: Cell Signaling (#9212), Mouse anti-FLAG: Sigma (F1804),Goat anti mouse IgG H&L (Alexa Fluor®647): abcam (ab150115), Goat anti sema3C: Santa-Cruz (sc-27796).

### Kits and reagents

NucleoSpin RNA Plus was from Macherey-Nagel (Cat. No. 740984.50). The qScript cDNA Synthesis Kit was from Quantabio (Cat. No. 95047-100). The WST-1 kit was from Roche Diagnostics (Cat. No. 05 015 944 001). The Senescence β-Galactosidase staining kit was from Cell Signaling technology (Cat. No. 9860) and Lipofectamine 3000 was purchased from Invitrogen (Cat. No. L3000008).

### Plasmids

The NSPI-CMV-MCS-myc-His lentiviral expression vector was kindly provided by Dr. Gal Akiri (Mount Sinai School of Medicine, NY) (Akiri *et al*., 2003). The plexin-A2/myc plasmid was kindly provided by Dr. Oded Behar (Hebrew University, Jerusalem, Israel). The pDonor221 and pLenti6/V5- DEST plasmids were purchased from Invitrogen. The pENTR1A-GFP-N2 (#19364) plasmid was from Addgene [deposited by Eric Campeau (Campeau *et al*., 2009)]. The pSpCas9(BB)-2A-GFP (#48138) plasmid was also from Addgene [deposited by Feng Zhang (Ran *et al*., 2013)]. The ΔNRF (pCMV dR 8.74) and pMD2-VSV-G vectors for lentivirus production were kindly provided by Dr. Tal Kafri (University of North Carolina at Chapel Hill, NC). The shRNA-containing lentiviral vectors were purchased from Sigma Aldrich.

### Cell lines

HUVECs were isolated and cultured as previously described (Gospodarowicz *et al*., 1978). HEK293 cells were purchased from the American Type Culture Collection (ATCC) and cultured as previously described (Kessler *et al*., 2004). HEK293-FT cells were purchased from Invitrogen. U87MG cells were Purchased from the ATCC and cultured as previously described (Sabag *et al*., 2014). U373MG, A172 and T98G cells were grown as described for U373MG cells (Sabag *et al*., 2012)(Sabag et al.). U118MG cells were cultured as previously described for HEK293 cells, the medium was also supplemented with 0.1mM non-essential amino acids and 1 mM sodium pyruvate (Biological Industries). All these cell lines were purchased from the ATCC. Puromycin, 2 µg/ml (Sigma) or blasticidin, 20 µg/ml (Invitrogen) were used to select infected cells.

### Inhibition of plexin expression with shRNA-expressing lentiviruses

Mission plasmids directing expression of shRNAs targeting PlexinA2 (TRCN0000061499 (ShPlexA2#1) and TRCN0000061501 (ShPlexA2#2)) or PlexinA4 (TRCN0000078683) were purchased from Sigma Aldrich. The production of the lentiviruses, infection of cells and the selection of shRNA expression cells in HEK293 cells were performed as described previously (Varshavsky *et al*., 2008).

### Generation of plexin-A2 knock-out cells using CRISPR/Cas9

U87MG cells were transfected with a pSpCas9(BB)-2A-GFP plasmids containing two different single guide RNAs (sgRNAs) sequences for plexin-A2 (Fig. S2A). The sgRNAs were chosen by using the CRISPR Design Tool (http://crispr.mit.edu/). Fluorescence-activated cell sorting (FACS) was performed 48hr after the transfection, for selection of the GFP expressing cells. The sorted cells were submitted to limiting dilution cloning. Clones containing mutations in the areas adjacent to the sgRNA target area were identified by PCR and the areas adjacent to the DNA region containing the sgRNA target sequence were then sequenced. The DNA sequences obtained were compared to the consensus DNA sequence of the gene and thoroughly examined in order to find the ones with insertion/deletion mutations causing frame shift disruption in both alleles (Cong *et al*., 2013) using an online tool (http://shinyapps.datacurators.nl/tide/) (Brinkman *et al*., 2014;Etard *et al*., 2017). Clones which were found to have null mutations in both alleles were submitted to western blot analysis for confirmation of the protein absence and later for cytoskeleton collapse experiments with sema3B.

### Quantitative real-time PCR

Quantitative real-time PCR was performed using a StepOne Plus Real Time PCR System with TaqMen Universal PCR Master Mix, according to the instructions of the manufacturer (Applied biosystems). The normalizing gene was RPLPO. Data was analyzed by the StepOne Software (Applied biosystems) using the relative Quantitation-Comparative C_T_ method. The following primers were used: PlexinA2-Hs00300697, PlexinA4-Hs00297356 and RPLPO-Hs99999902.

### Expression and production of semaphorins

Class 3 semaphorin cDNAs were subcloned into the NSPI-CMV-myc-his lentiviral expression vector as described (Varshavsky *et al*., 2008;Kigel *et al*., 2008). The production of lentiviruses and the generation of conditioned media containing various semaphorins were performed as described (Kigel *et al*., 2008).

### Cytoskeletal collapse experiments

Cytoskeleton collapse assays using U87MG cells were performed as previously described (Kigel *et al*., 2011;Sabag *et al*., 2014) using HEK293 cell-derived conditioned medium containing recombinant sema3B or control conditioned medium from cells containing empty expression vectors. To stabilize pH we also added HEPES buffer (10 mM, pH 7.2). Cells were photographed after a 30 min incubation in a humidified incubator at 37°C using a phase-contrast inverted microscope.

### Expression of recombinant plexins

The full-length cDNA of plexin-A2 was cloned into the gateway pDonor221 plasmid and then transferred by recombination into the pLenti6/V5-DEST lentiviral expression vector in frame with a C-terminal V5 tag according to the instructions of the manufacturer (Invitrogen), as previously described (Sabag *et al*., 2014). A deletion mutant of plexin-A2 containing the extracellular and transmembrane domains (amino acids 1–1323) of the protein fused to a C-terminal FLAG tag (A2ExTm) was cloned into the gateway entry vector, pENTR1A-GFP-N2, and then transferred by recombination into the pLenti6/V5-DEST lentiviral expression vector. A deletion mutant of plexin-A2 containing the intracellular and transmembrane domains amino acids (1214–1894) of the protein (A2InTm) was generated by truncation of bases 3640-5685 of the plexin-A2 cDNA. The truncated plexin-A2 cDNA was then assembled using NEBuilder HiFi DNA Assembly Master Mix, according to the instructions of the manufacturer (New England Biolabs) into the NSPI-CMV-MCS-myc-His lentiviral expression vector in frame with an N-terminal signal peptide of human plexin-A4 and a V5 tag. A2InTm was subsequently subcloned similarly into the pLenti6/V5-DEST lentiviral expression vector. Introduction of point mutations into the cDNA of A2InTm at the Rho GTPase binding domain (RBD domain) was generated by replacement of the LVP motif with GGA following a previous publication (Fig. S6A) (Liu and Strittmatter, 2001). Introduction of point mutations into the cDNA of A2InTm at the FERM domain binding site was generated by replacement of the KRK motif with a triplet of alanine following a previous publication (Toyofuku *et al*., 2005) (Fig. S6A). Both of these mutants of A2InTm were similarly cloned into the pLenti6/V5-DEST lentiviral expression vector. We also received a plexin-A2 cDNA variant containing a double point mutant at the FYN dependent phosphorylation sites of plexin-A2 (Y1605F/Y1677F) which was kindly provided by Dr. Riley M. St. Clair (University of Vermont, Burlington, VT) (St Clair *et al*., 2017). We then generated A2InTm mutated at these FYN dependent phosphorylation sites by truncation of bases 3640-5685 of the plexin-A2 cDNA variant and similarly cloning it into the pLenti6/V5-DEST lentiviral expression vector. Additionally, introduction of point mutations into the cDNA of A2InTm at the cytosolic juxtamembrane dimerization interface was generated by replacement of methionine with phenylalanine following a previous publication (Barton *et al*., 2015) (Fig. S6A) and similarly cloning it into the pLenti6/V5-DEST lentiviral expression vector. Production of lentiviruses using these plasmids and stable infection of target cells were performed essentially as previously described (Varshavsky *et al*., 2008). Target cells expressing plexin-A2 or a deletion mutant of plexin-A2 containing the extracellular and transmembrane domains, were isolated from the pool of the infected cells using FACS. U87MG KO clone 54 cells in which we expressed the cDNA encoding the full length plexin-A2 cDNA (2.10^7^ cells) or U87MG cells in which we expressed cDNA encoding a dominant-negative truncated plexin-A2 lacking the intracellular domain of plexin-A2 (1.10^7^ cells) were washed with PBS and trypsinized, then resuspended in phosphate-buffer saline (PBS) with 0.2% bovine serum albumin (BSA) and with 10mM EDTA then incubated with Goat anti plexin-A2-APC (RαD systems) for 40 min in the dark at 25 °C for staining plexin-A2 or with Mouse anti-FLAG (Sigma) for 40 min at 4°C followed by Goat anti mouse IgG H&L (Alexa Fluor®647)(abcam) for 30 min in the dark at 4 °C for staining dominant-negative truncated plexin-A2. After staining, cells were sorted using a BD FACSAria IIIu cell sorter.

### Proliferation assays

U87MG cells (1×10^4^ cells/well) or HUVEC (2×10^4^ cells/well) were cultured as previously described (Tessler and Neufeld, 1990;Sabag *et al*., 2014). Proliferation assays were conducted, and the numbers of adherent cells determined using either a coulter-counter as previously described (Kigel *et al*., 2011) or using the WST-1 assay in which case 3×10^3^ cells were seeded per well. Te assay was performed according to the instructions of the vendor (Roche). The population doubling time was computed as the ln(2)/slope of the proliferation growth curve using Prism software. For clonal colony formation assay, U87MG cells (300 cells/well) were seeded in complete growth medium (10% FCS) and allowed to form colonies in 16 days. Colonies were fixed with PFA (4.0% v/v), stained with crystal violet (0.5% w/v) and photographed using an ImageQuant LAS-4000 machine.

### Soft-agar colony formation assay

Soft agar colony formation from U87MG cells was performed as previously described (Kigel *et al*., 2008). The number of colonies and their average area were determined using the Image-pro premier morphometric software.

### Immunofluorescence

For the staining of the cytoskeletal components, U87MG cells were plated on glass coverslips coated with fibronectin and actin fibers visualized as previously described (Guttmann-Raviv *et al*., 2007).

#### Senescence associated β-Galactosidase assay

The staining of SA-β galactosidase in cells at pH-6 was performed using a senescence β-Galactosidase staining kit according to the instructions of the manufacturer (Cell signaling). Measurement of the absorbance at 420 nm from SA-β galactosidase activity detectable at pH 6.0 was performed as described (Lee *et al*., 2006).

### Cell cycle analysis

Cultured cells (1.5.10^6^ cells) were washed with PBS and trypsinized, then fixed in 70% ethanol at -20 °C for overnight. The cells were then incubated in a solution containing 0.1 mg/ml RNase (Thermo Fisher Scientific) and 40 μg/ml propidium iodide (Sigma) at room temperature for 30 min, and finally cell cycle analysis was performed using flow cytometry (LSRFortessa; BD Biosciences) and FCS Express Flow Cytometry Software.

### Phosphorylation assays

The phosphorylation of p38 MAPK and AKT were determined as previously described for ERK1/2 phosphorylation (Guttmann-Raviv *et al*., 2007) using western blot analysis. Quantification of the stained bands intensity was done using an ImageQuant Las-4000 machine.

#### Tumor formation assays

U87MG control cells or cells that were silenced for plexin-A2 were washed, suspended in 100 µL PBS, subcutaneously injected into 4 week old Athymic/Nude mice (2×10^6^ cells/mouse) and the development of tumors was measured twice a week using calipers. At the end of the experiment the tumors were excised and weighed. All the animal experiments were approved by the Technion ethics committee.

### Statistical analysis

Statistical significance was determined in most cases using the Mann-Whitney one-tailed non-parametric test or by using the one-way ANOVA test followed by Bonferroni’s multiple comparison post-test, unless otherwise stated. Cell proliferation experiments were performed in quadruplicate and the variation between quadruplicates did not exceed 10%. Error bars represent the standard error of the mean. The following designations were used in the figures: *P<0.05, **P<0.01 and ***P<0.001. All experiments were repeated independently at least three times unless otherwise stated.

### Ethics statement

All the authors of this article have given their informed consent to the article. The animal studies were all conducted according to the NIH guidelines and were approved by the Institutional Review Board of the Technion.

## ACKNOWLEDGMENTS

This work was supported by a grant from the Israel Science Foundation (ISF) to GN.

---

**Supp. Figure 1.**
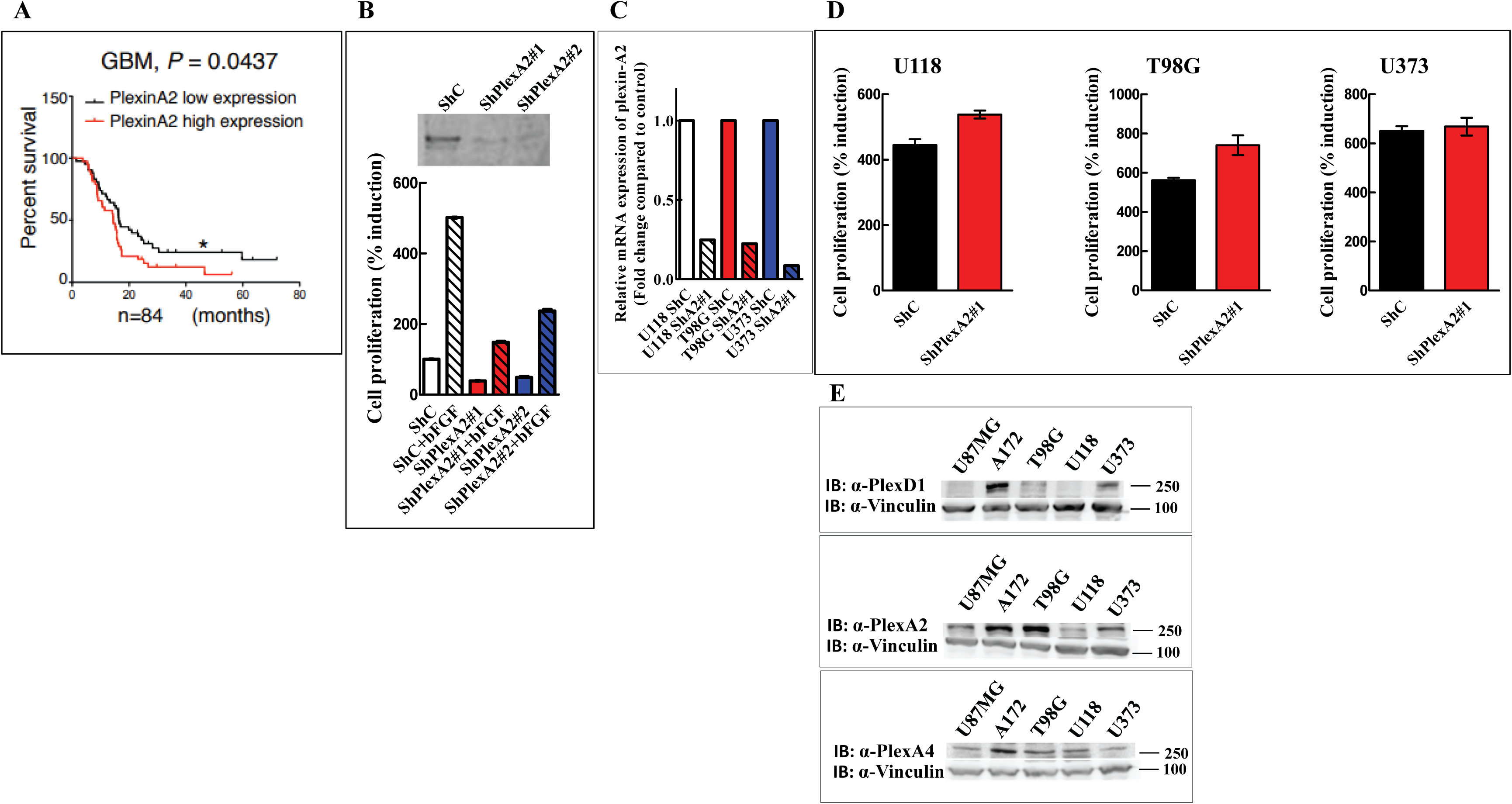
Expression levels of plexin-A2 in human glioma and the effect of silencing plexin-A2 in HUVEC cells or in additional glioblastoma multiforme cells on their proliferation: **(A)** Correlation of PlexinA2 mRNA expression with survival of GBM patients. Murat brain database, Oncomine (Murat *et al*., 2008;Man *et al*., 2014) **(B)** Effects of two shRNA species targeting plexin-A2 on the proliferation of human umbilical vein derived endothelial (HUVEC) cells. HUVEC cells expressing a non-specific shRNA (ShC) or HUVEC cells in which the expression of plexin-A2 was silenced using two shRNAs (ShPlexA2#1, or ShPlexA2#2) were seeded in triplicates (2×10^4^ cells/well) in 24 well plates in the presence or absence of bFGF (5 ng/ml). Adherent cells were counted in coulter counter after 3 days. Shown is the percentage of cells in day 3 as compared to the number of cells expressing a control shRNA (ShC) taken as 100%. The experiment was repeated twice with similar results. Error bars represent the standard error of the mean. **(C)** Lentiviruses were used to express control (ShC) or a plexin-A2-targeting shRNA ShplexA2#1 (ShA2#1) in U118, T98G and U373 cells. The expression of plexin-A2 was then examined using qRT-PCR. **(D)** U118, T98G and U373 cells were infected with lentiviruses encoding a control shRNA (ShC) or a shRNA targeting plexinA2 (ShPlexA2#1). Each group of cells was seeded in quadruplicate in 96 well dishes (3×10^3^ cells/well). Cell proliferation was measured using the WST-1 proliferation assay as described in materials and methods, and the values presented were calculated as described in Fig 1B. Error bars represent the standard error of the mean. **(E)** Western blots prepared from cell lysates of U87MG, A172, T98G, U118 and U373 glioblastoma multiforme cells were probed with antibodies directed against plexin-D1, plexin-A2 or plexin-A4. Antibodies directed against vinculin were used to compare loading.

**Supp. Figure 2.**
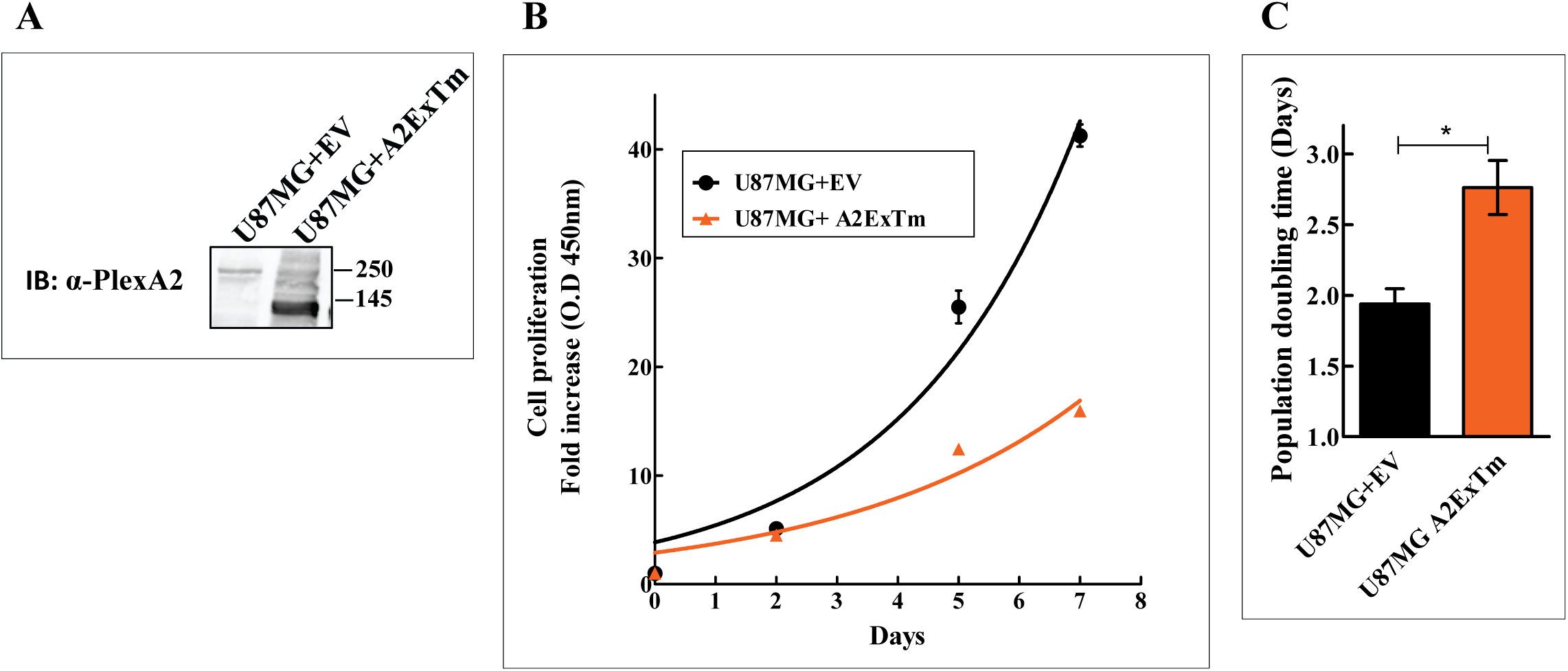
A dominant negative form of plexin-A2 lacking the intracellular domain (A2ExTm) inhibits the proliferation of U87MG cells: **(A)** U87MG cells were infected with an empty lentiviral expression vector (U87MG+EV) or with a lentiviral vector directing expression of A2ExTm (U87MG+A2ExTm). Western blots prepared from cell lysates were probed with an antibody directed against plexin-A2. **(B)** U87MG cells infected with an empty lentiviral expression vector (EV) or with a lentiviral vector directing expression of A2ExTm were seeded in 96 well dishes (3×10^3^ cells/well). Cell proliferation was measured using the WST-1 proliferation assay as described in methods. **(C)** The average population doubling time of the cells, from three independent experiments such as the one shown in B. Statistical significance was evaluated using the one tailed Mann-Whitney test. Error bars represent the standard error of the mean.

**Supp. Figure 3.**
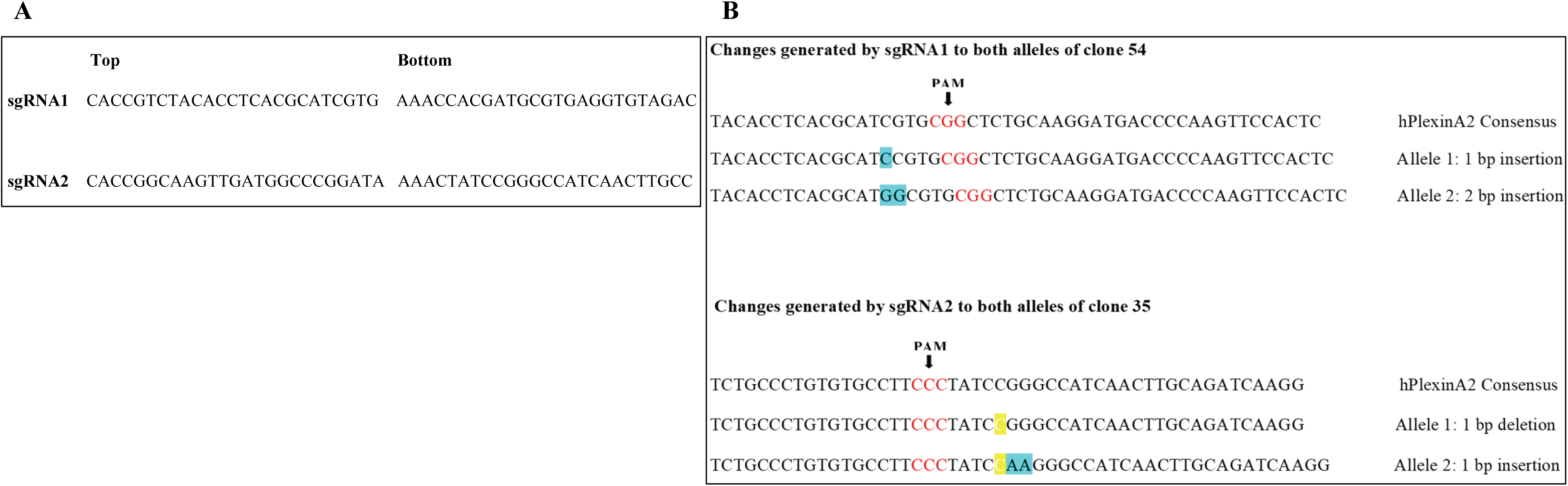
Guide RNAs used for the generation of plexin-A2 knock-out U87MG cells: **(A)** Sequences of the sgRNAs encoding primers used for CRISPR/Cas9 mediated knock-out of the plexin-A2 gene in U87MG cells. **(B)** Sequence analysis of two plexin-A2 knock-out U87MG clones of cells. Shown are the target sequences in the plexin-A2 gene and the frame shift mutations that were introduced into the two alleles of the plexin-A2 gene in knock-out clones 54 and 35 which were determined by sequencing and analyzed as described in materials and methods.

**Supp. Figure 4.**
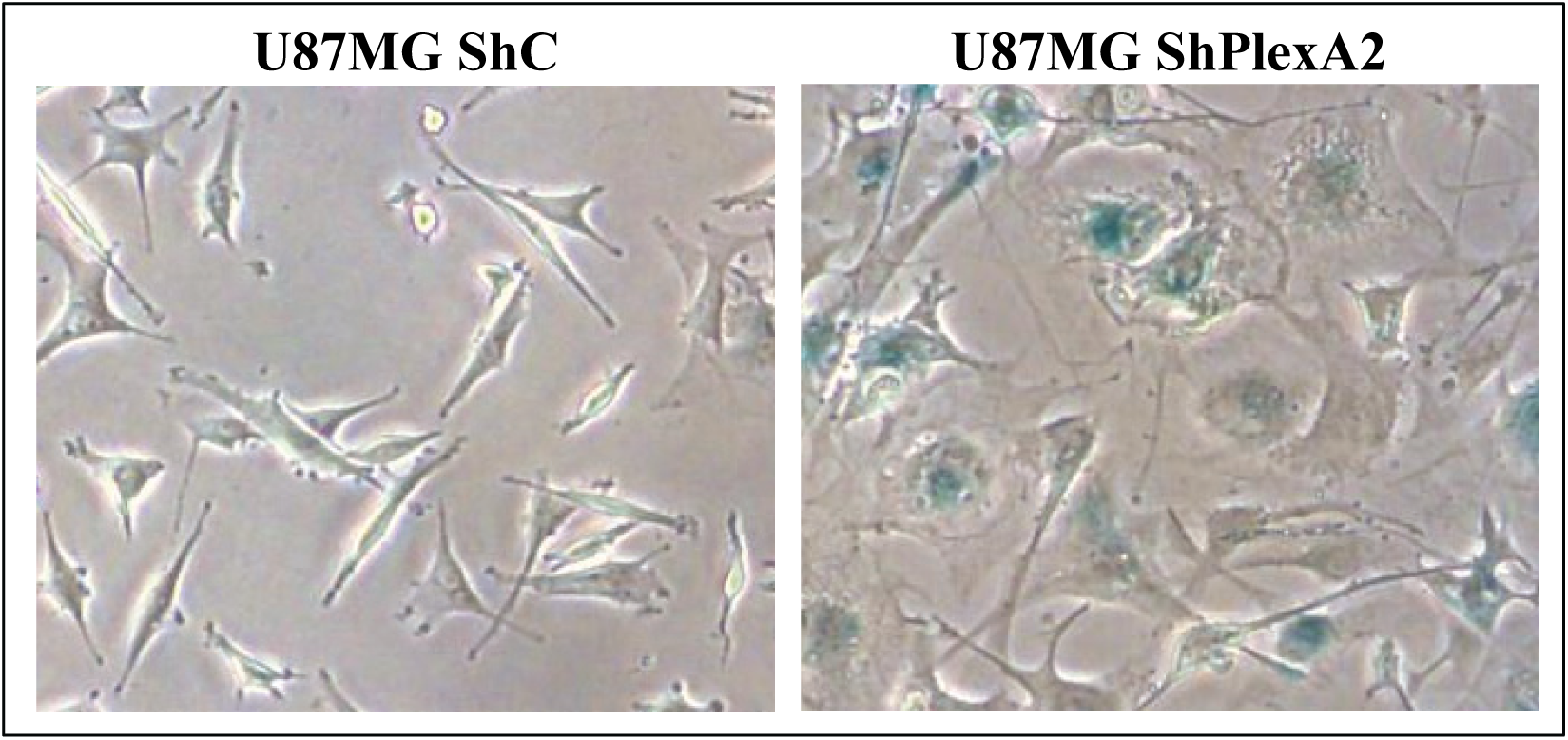
U87MG cells silenced for plexin-A2 expression acquire some properties of senescent cells: U87MG cells expressing a non-specific shRNA (ShC) or U87MG cells in which the expression of plexin-A2 was silenced using a shRNA (ShPlexA2#1) were assayed at pH-6 for the expression of the senescence marker SA-β galactosidase.

**Supp. Figure 5.**
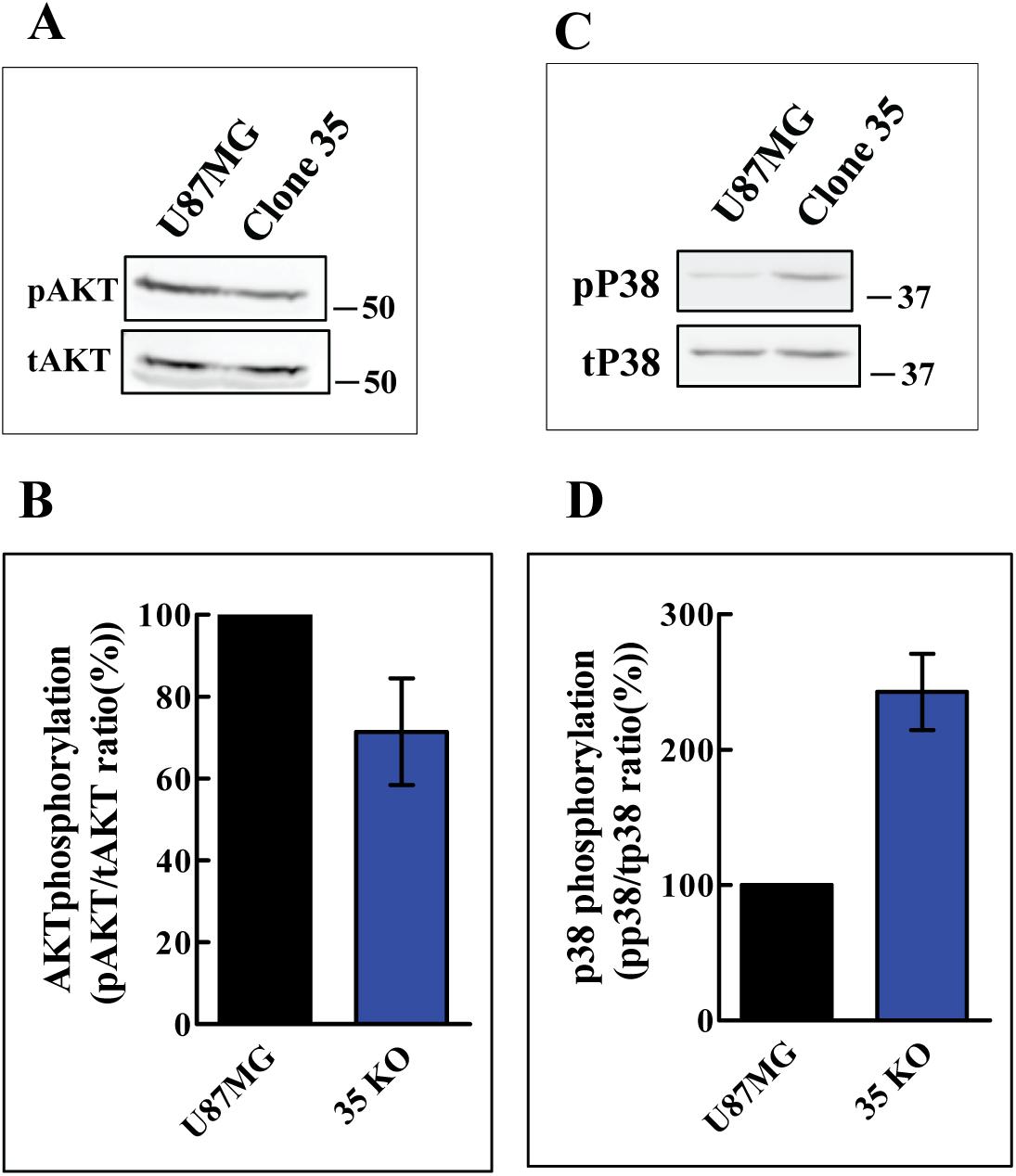
The phosphorylation of AKT is inhibited and that of p38 induced in clone 35 plexin-A2 knock-out cells: **(A)** The phosphorylation levels of AKT were assayed using western blot analysis of cell lysates and an antibody directed against phosphorylated AKT (ser473). Loading was assessed using an antibody directed against total AKT. Shown is a representative western blot. **(B)** The effect of plexin-A2 knock-out on the average phosphorylation level of AKT was determined in 4 independent experiments as described in methods. Shown is the average phosphorylation level of AKT. Error bars represent the standard error of the mean. **(C)** P38 phosphorylation levels were assayed using western blot analysis of cell lysates and an antibody directed against phosphorylated p38 (Thr180/Tyr182). Loading was assessed using an antibody directed against total p38. Shown is a representative experiment. **(D)** The effect of plexin-A2 knock-out on the average phosphorylation level of p38 derived from 4 independent experiments is shown. Error bars represent the standard error of the mean.

**Supp. Figure 6.**
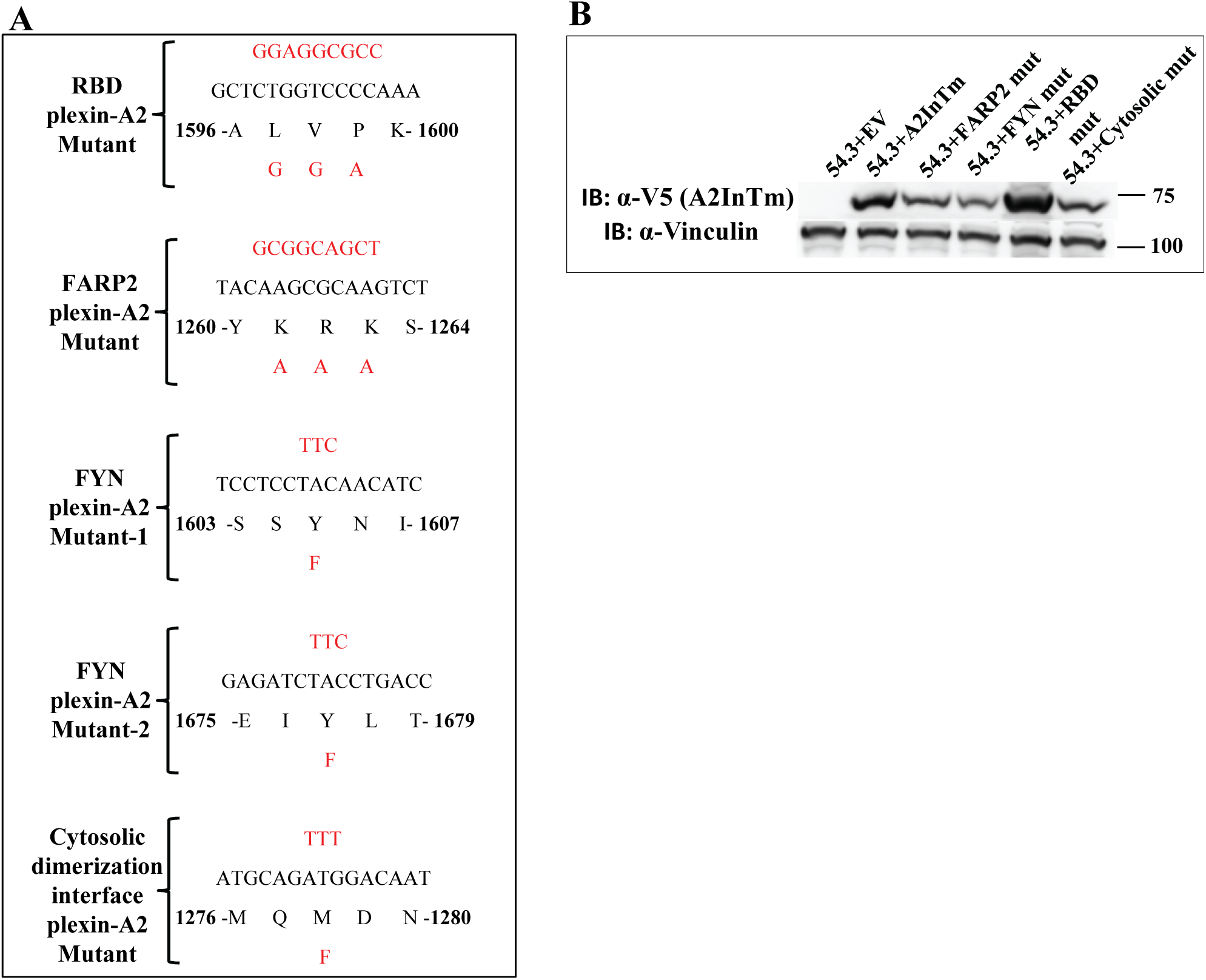
Generation of A2InTm variants containing point mutations: **(A)** Sequences of point mutations introduced into the A2InTm cDNA encoding the intracellular and trans-membrane domain of plexin-A2 **(B)** Western blots prepared from cell lysates of clone 54.3 knock-out cells (54.3) expressing cDNAs encoding an empty expression vector (EV), a V5 tagged A2InTm or A2InTm variants containing point mutations in the indicated subdomains of the plexin-A2 intracellular domain were probed with antibodies directed against the V5 epitope tag of A2InTm or vinculin.

